# Y chromosome linked UTY modulates sex differences in valvular fibroblast methylation in response to nanoscale extracellular matrix cues

**DOI:** 10.1101/2024.05.13.593760

**Authors:** Rayyan M. Gorashi, Talia Baddour, Sarah J. Chittle, Nicole E. Félix Vélez, Michaela A. Wenning, Kristi S. Anseth, Luisa Mestroni, Brisa Peña, Peng Guo, Brian A. Aguado

## Abstract

Aortic valve stenosis (AVS) is a progressive disease wherein males more often develop valve calcification relative to females that develop valve fibrosis. Valvular interstitial cells (VICs) aberrantly activate to myofibroblasts during AVS, driving the fibrotic valve phenotype in females. Myofibroblasts further differentiate into osteoblast-like cells and produce calcium nanoparticles, driving valve calcification in males. We hypothesized the lysine demethylase UTY (ubiquitously transcribed tetratricopeptide repeat containing, Y-linked) decreases methylation uniquely in male VICs responding to nanoscale extracellular matrix cues to promote an osteoblast-like cell phenotype. Here, we describe a hydrogel biomaterial cell culture platform to interrogate how nanoscale cues modulate sex-specific methylation states in VICs activating to myofibroblasts and osteoblast-like cells. We found UTY modulates the osteoblast-like cell phenotype in response to nanoscale cues uniquely in male VICs. Overall, we reveal a novel role of UTY in the regulation of calcification processes in males during AVS progression.

## Introduction

Aortic valve stenosis (AVS) is sexually dimorphic disease where aortic valve tissue progressively undergoes fibro-calcification^1^. AVS impacts approximately 13% of the population aged over 75, and gender-based disparities exist with clinical management of AVS, given retrospective studies suggesting women are under recommended for valve replacements^2^. Given that valve replacement patients are also at risk for increased co-morbidities after the procedure, a significant need exists to identify non-surgical based strategies to delay the onset of AVS^3^. Clinical research further suggests the degree of fibro-calcification in the valve depends on biological sex, where males exhibit increased calcification relative to females^4,5^. Given the sex dimorphisms observed in valve tissue during AVS onset, biological sex must be considered when identifying paths toward non-surgical pharmacologic treatments for AVS.

Our collective understanding of the sex-specific fibro-calcification processes in valve tissue during AVS remain limited. Valvular interstitial cells (VICs) are the resident fibroblast-like cells in valve tissue that are known to regulate valve fibro-calcification^4^. In early stages of AVS, VICs chronically activate to myofibroblasts and express alpha smooth muscle actin (aSMA) stress fibers^6^. In moderate to late stages of AVS, myofibroblasts further differentiate into osteoblast-like valve cells^7,8^. Osteoblast-like valve cells are characterized by the nuclear localization of runt-related transcription factor (RUNX2) and contribute to the accumulation of stiff, spherical calcium phosphate nanoparticles in valve tissue^9,10^. Over time, nanoparticles increase in size and abundance and further exacerbate valve fibro-calcification^4^. We posit the initiation of sex specific AVS progression partially depends on VIC interactions with the valve extracellular microenvironment.

Calcific lesions observed in healthy and late stage AVS tissue contain dense, mineralized spherical nanoparticles (NPs) of varying size in the valve extracellular microenvironment^11^. Prior studies have shown that osteoblast-like cells experience altered cytoskeletal dynamics after culture on substrates containing stiff NPs^12^. In particular, the NPs affected actin organization, disrupted microtubule networks, and reduced overall cell spreading. VICs interact with their surrounding microenvironment via transmembrane integrin receptors^13,14^. Integrins are directly linked to the actin cytoskeleton and mediate cell response to external stimuli through alterations to the nuclear chromatin structure^15–17^. Prior studies have also identified differences in chromatin structure of VICs from both healthy and diseased patients^18,19^. As such, chromatin re-arrangement in male and female VICs in response to nanoscale cues in the extracellular matrix is hypothesized to modulate sex-specific cellular phenotypes during valve fibro-calcification.

Furthermore, cellular responses to the extracellular microenvironment also depend on intracellular sex chromosome effects^20–23^. Sex differences in AVS progression have been attributed to both patient-specific sex hormone levels and sex chromosome activity^24–26^. To detangle the effects of sex hormones from sex chromosome activity, approaches using hydrogel biomaterials to recapitulate the valve extracellular matrix have been used to show female VICs have increased myofibroblast activation due to genes that escape X chromosome inactivation^27,28^. Outside of VICs, previous work has also shown hydrogels to be critical tools to understand sex-specific mechanotransduction in endothelial cells cultured *in vitro*^29,30^. Our understanding of how VICs respond to nanoscale cues in the extracellular matrix will continue to harness *in vitro* culture tools to probe sex-specific cell-nanomaterial interactions and subsequent sex-specific effects on cell phenotypes.

Here, we interrogated how nanoscale matrix cues modulate the VIC-to-myofibroblast-to-osteoblast transition in males and females. We first performed single cell sequencing on human valve tissue samples from AVS patients, revealing upregulation of pro-osteogenic genes and decreased expression of the Y-chromosome linked histone demethylase, known as Ubiquitously Transcribed Tetratricopeptide Repeat Containing, Y-linked (UTY) in male VICs. Next, we engineered a bioinspired hydrogel cell culture platform to probe how nanoscale matrix cues modulate VIC phenotypes via UTY. We have previously used hydrogel biomaterials to culture primary VICs isolated from aortic valve leaflets and determined that sex chromosome linked genes modulate sex differences in VICs phenotypes^27,28^. In our current approach, we incorporated stiff, polystyrene nanoparticles (PS-NPs) into a water-swollen crosslinked poly(ethylene glycol) (PEG) norbornene hydrogel to mimic nanoscale matrix cues in aortic valve tissue. After culture on our PS-NP hydrogel, our results collectively show male-specific osteoblast-like responses to nanoscale stiffness cues. After *UTY* knockdown, we reveal upregulation of genes involved in osteogenic differentiation, ultimately promoting calcification processes in VICs. We further validated our bioinspired hydrogel platform by comparing the *in vitro* male VIC transcriptome on PS-NP hydrogels with our single cell transcriptomic dataset from human tissues. Overall, we have potentiated a role of the Y-linked UTY lysine demethylase in causing sex-specific effects that guide male VICs towards calcification in AVS.

## Results

### Single-cell sequencing reveals distinct VIC populations in diseased human aortic valves

We first characterized sex-specific valvular interstitial cell (VIC) populations in diseased aortic valves relative to healthy valves. We conducted our sex-separated analysis by combining publicly available AVS patient data and our own patient samples (**Supplementary Table 1**). After clustering, our uniform manifold approximation and projection (UMAP) plots show endothelial cells, macrophages, T-cells, and 5 distinct VIC populations, in both male and female patients (**Figure 1A**, **Supplementary Figure 1A**). Cell populations were determined using conserved biomarkers and further identified by distinct biomarkers (**Supplementary 1B-C**, **Supplementary Figures 2 and 3**). In both male and female patients, VIC population percentages significantly increase with disease and comprises the majority of detected cells in the aortic valve tissue (**Figure 1B**, **Supplementary Figure 4A-B**).

**Figure 1.**
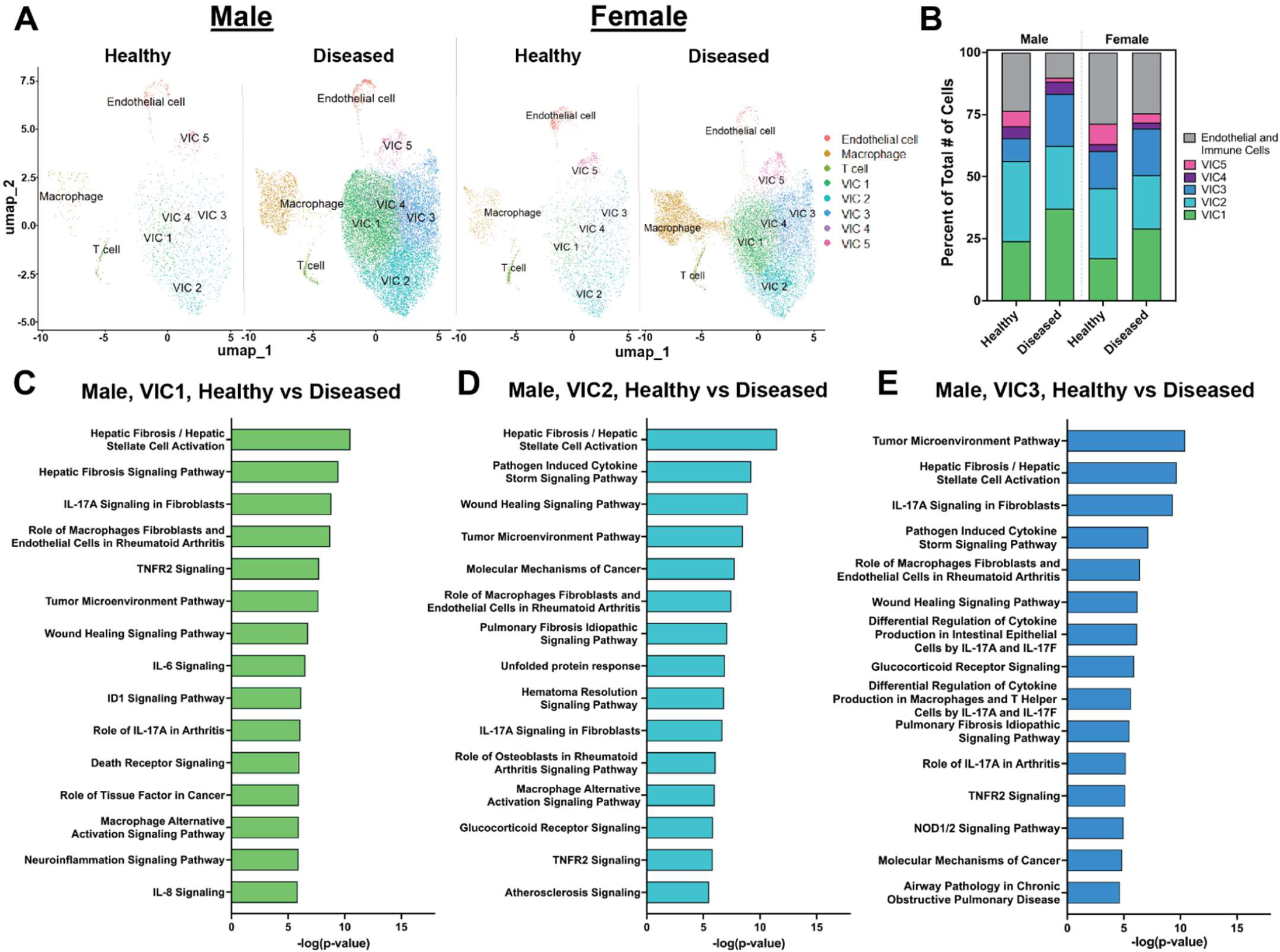
Single-cell sequencing and sex-separated analysis of human AVS patients reveal clustering of VIC populations. (**A**) UMAP clustering and cell population distribution in healthy and diseased patient samples for males and females. (**B**) Distribution of VIC and non-VIC population percentages in healthy and diseased patient samples for males and females. Ingenuity pathway analysis for male (**C**) VIC1, (**D**) VIC2, and (**E**) VIC3.

Ingenuity pathway analysis revealed significant associations with pathways related to pro-inflammatory and pro-fibrotic pathways, senescence, and interleukin-17 signaling for the VIC1 population (**Figure 1C**). The VIC2 and VIC3 populations showed significant associations to pathways such as interleukin-17 signaling, tumor necrosis factor receptor-2 signaling, chondrocytes in inflammatory diseases, and toll-like receptor signaling (**Figure 1D-E**). VIC4 and VIC5 showed significant associations to pathways involving senescence, death receptor signaling, and apoptosis signaling (**Supplementary Figure 5**).

### Hydrogels containing polystyrene nanoparticles mimic the valve microenvironment

We next sought to engineer a hydrogel cell culture platform that recapitulates the valve matrix that contains nano-scale calcium phosphate particles. To determine the appropriate particle size scale, we characterized the nanoscale matrix features from age-matched male and female aortic valve tissue from two healthy patients and two patients diagnosed with AVS. Using scanning electron microscopy (SEM), we visualized the accumulation of nano-scale calcium phosphate particles in aortic valve tissue from male and female AVS patients relative to healthy patients (**Figure 2A**). After quantifying particle diameter, we observe that calcium phosphate particles in male diseased patients have the highest frequency distribution between 100-2000 nm. The largest particles from our severely stenotic male AVS patient sample measured 10000 nm. In female diseased patients, we observe the highest frequency distribution between 50-1000 nm. In our severely stenotic female AVS patient sample, the largest particles measured 21000 nm (**Supplementary Figure 6A-C**).

**Figure 2.**
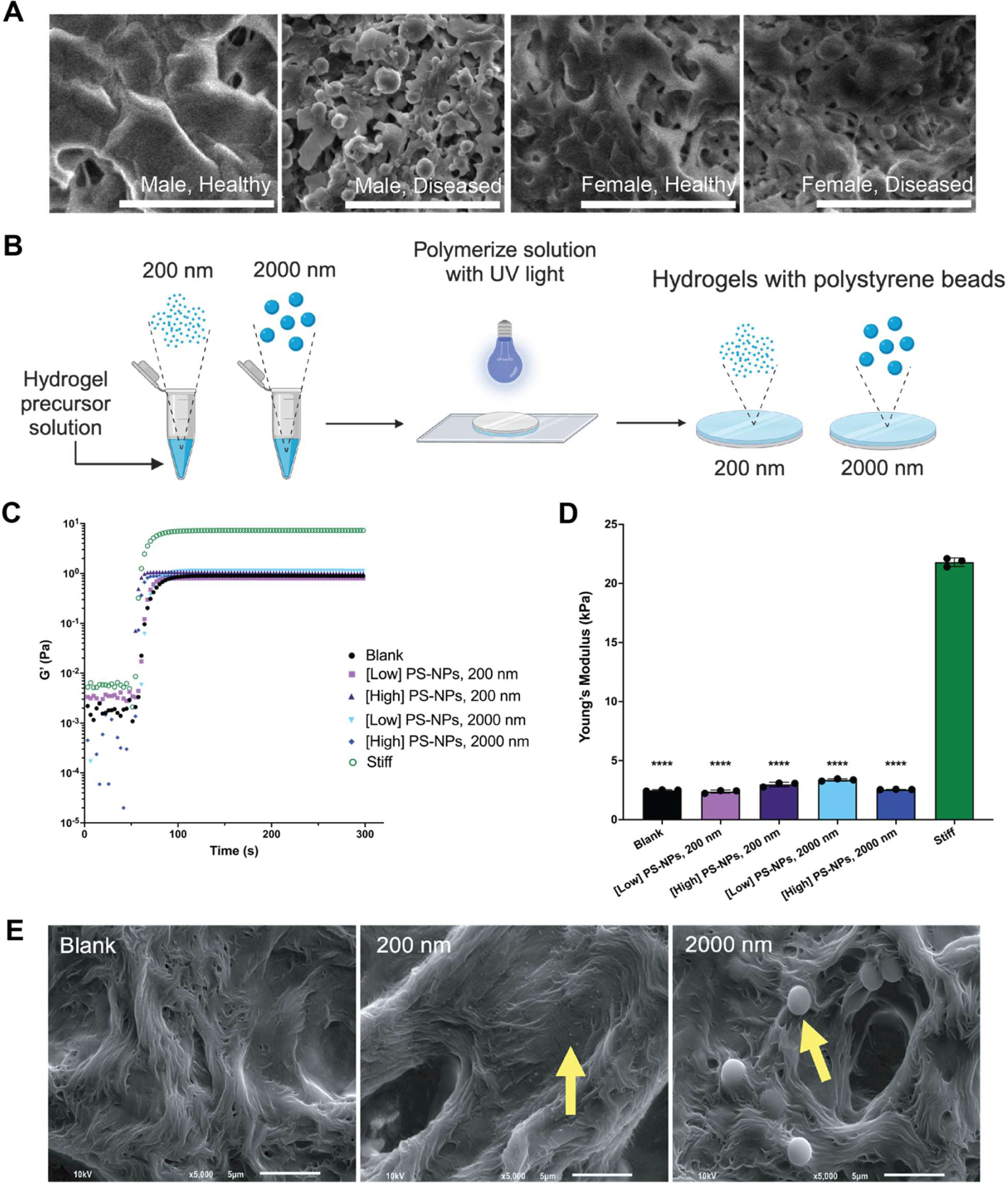
PS-NP hydrogels recapitulate nano-scale features in aortic valve tissue ECM. (**A**) Scanning electron microscopy (SEM) images of male healthy, diseased, and female diseased human aortic valve leaflets. Scale bar = 5 μm. (**B**) Schematic of process to incorporate polystyrene nanoparticles (PS-NPs) into PEG-Norbornene hydrogel precursor solutions to create PS-NP hydrogels adhered to glass coverslips via UV polymerization. Figure drawn with BioRender. (**C**) Rheological characterization of UV-mediated polymerization of soft, stiff, and PS-NP hydrogels made with 200 nm and 2000 nm nanoparticles (n=3 representative measurements) at low and high concentrations (0.02 mg/ml and 0.7 mg/ml, respectively). (**D**) Young’s modulus values for soft, stiff, and PS-NP hydrogels (N=3 gels). Significance determined via one-way ANOVA (****p<0.0001 denoting statistical significance between hydrogel formulations). Mean ± SD shown. (**E**) SEM of blank and PS-NP hydrogels (200 nm, 2000 nm). Yellow arrows denote PS-NPs. Scale bar = 5 μm.

To mimic the range of observed particle sizes in the valve tissue microenvironment, we incorporated 200 or 2000 nm diameter polystyrene nanoparticles (PS-NPs) into our poly(ethylene glycol) (PEG) hydrogel network to create a matrix containing nano-scale particles (**Figure 2B**). We quantified hydrogel storage and elastic modulus using shear rheology (**Figure 2C-D**) and surface stiffness using atomic force microscopy (**Supplementary Figure 7).** We determined that the elastic modulus (E) of our soft, blank hydrogel matrix formulation (E = 2.48 ± 0.05 kPa) does not significantly change with the incorporation of 200 nm PS-NPs (E_200 nm PS-NPs, low_ = 2.37 ± 0.12 kPa and E_200 nm PS-NPs, high_ = 2.98 ± 0.19 kPa) and 2000 nm PS-NPs (E_2000 nm PS-NPs, low_ = 3.36 ± 0.09 kPa and E_2000 nm PS-NPs, high_ = 2.55 ± 0.02 kPa), relative to our stiff hydrogel formulation (E = 21.80 ± 0.36 kPa). We also utilized SEM to confirm PS-NP presence on the surface of the hydrogel matrix (**Figure 2E**).

### Sex-specific myofibroblast activation and osteoblast-like differentiation on nanoparticle hydrogels

To test the effect of PS-NP nanoparticles on sex-specific valve cell phenotypes, we isolated valvular interstitial cells (VICs) from porcine aortic valve tissue for culture on our PS-NP hydrogels (**Figure 3A**). First, using alpha smooth muscle actin (αSMA) stress fibers as a marker of myofibroblast activation, we observed female VICs having significantly higher levels of myofibroblast activation on both the blank hydrogel and PS-NP hydrogel conditions, compared to male VICs (**Figure 3B-C**). Second, we evaluated the nuclear localization of the transcription factor runt-related transcription factor 2 (RUNX2) as a marker of osteoblast-like differentiation. Male VICs displayed significantly higher levels of Runx2 nuclear localization on the blank hydrogel and most PS-NP hydrogel conditions, compared to female VICs (**Figure 3D-E**). Collectively, our results provide evidence that male and female VICs exhibit sex-specific differences in myofibroblast and osteoblast-like phenotypes in response to PS-NPs in culture. Though we detected these sex-specific phenotypes in response to PS-NP hydrogels, we did not observe consistent dose-dependent or PS-NP size-dependent trends.

**Figure 3.**
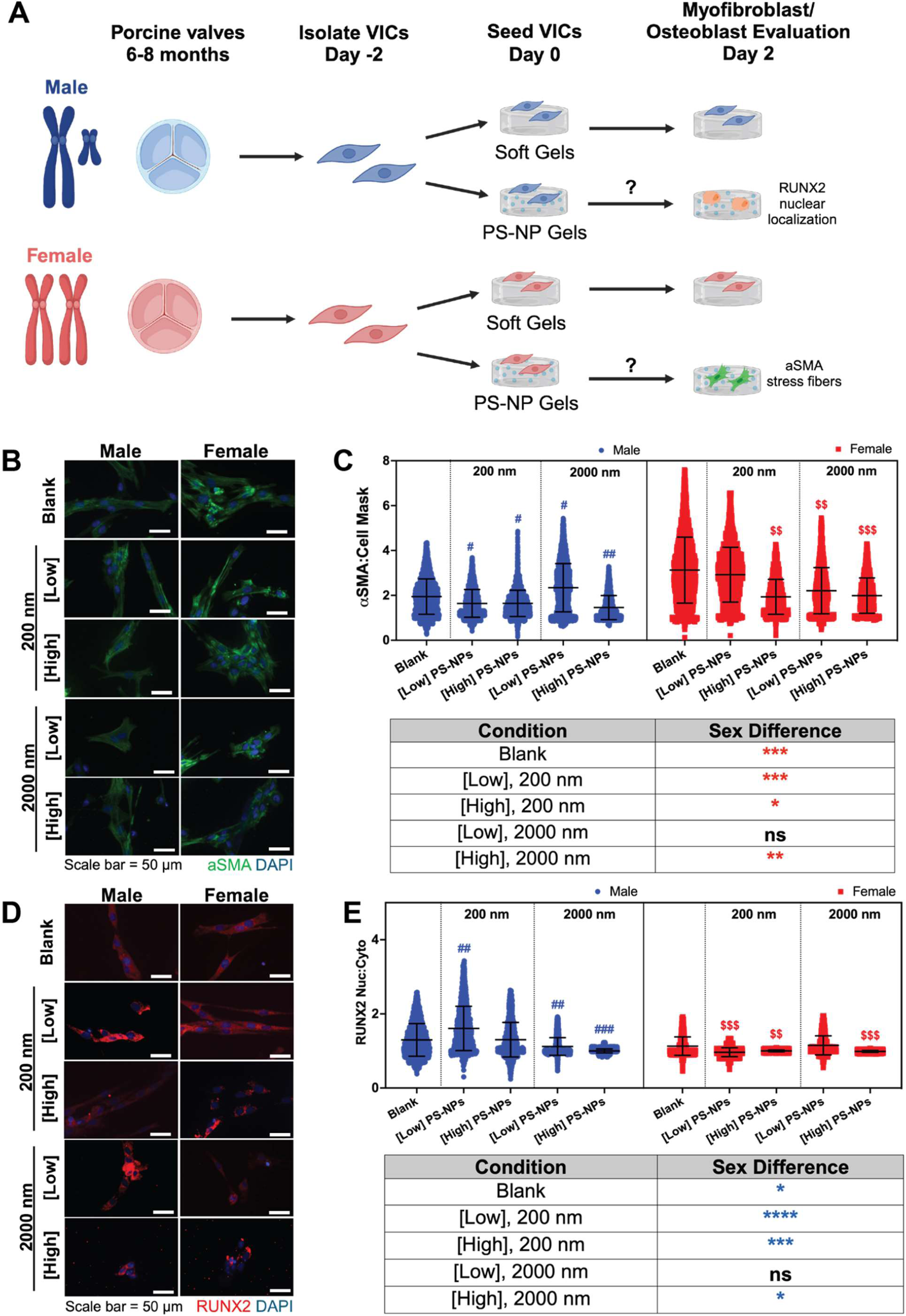
Male and female VICs exhibit sex-specific myofibroblast and osteoblast-like phenotypes. (**A**) Schematic describing experimental workflow for probing myofibroblast activation with α-smooth muscle actin (α-SMA) and osteoblast-like differentiation with runt related (RUNX2). Figure drawn with BioRender. (**B**) Representative confocal microscopy images of alpha smooth muscle stress fibers (green) and nuclei (blue). Scale bar = 50 μm. (**C**) Normalized α-SMA expression ratios for male and female VICs cultured on blank hydrogels with no PS-NPs, 200 nm PS-NPs, or 2000 nm PS-NPs at low and high concentrations (0.02 mg/ml and 0.7 mg/ml, respectively). Significance determined via Cohen’s d-test (*d < 0.2, **d < 0.5, ***d<0.8, ****d<1.4 denoting significance between biological sex in the provided table, red asterisk color indicates significant increase in female groups, ^#^d < 0.2, ^##^d < 0.5, ^###^d<0.8, ^####^d<1.4 denoting significance relative to male blank with no PS-NPs, ^$^d < 0.2, ^$$^d < 0.5, ^$$$^d<0.8, ^$$$$^d<1.4 denoting significance relative to female blank with no PS-NPs). (**D**) Representative confocal microscopy images of RUNX2 expression (red) and nuclei (blue). Scale bar = 50 μm. (**E**) RUNX2 nuclear to cytoplasmic intensity ratios of male and female VICs cultured on blank hydrogels with no PS-NPs, 200 nm PS-NPs, or 2000 nm PS-NPs at low and high concentrations (0.02 mg/ml and 0.7 mg/ml, respectively). Significance determined via Cohen’s d-test (*d < 0.2, **d < 0.5, ***d<0.8, ****d<1.4 denoting significance between biological sex in the provided table, blue asterisk color indicates significant increase in male groups, ^#^d < 0.2, ^##^d < 0.5, ^###^d<0.8, ^####^d<1.4 denoting significance relative to male blank with no particles, ^$^d < 0.2,

### Sex-specific methylation and acetylation states on PS-NP hydrogels

Recognizing that epigenetic modifications drive cellular phenotype plasticity via alterations to methylation and acetylation states in the nucleus^31^, we sought to characterize sex-specific epigenetic states in VICs on our PS-NP gels (**Figure 4A**). To understand cell-matrix interactions that modulate transcriptional activity, we sought to investigate methylation states in VICs cultured on PS-NP hydrogels. Male VICs displayed reduced tri-methylation across most hydrogel conditions relative to female VICs (**Figure 4B-C**). Interestingly, we observed dose-dependent and PS-NP size-dependent changes in male VIC methylation after culture on PS-NP hydrogels. Next, we determined acetylation states in VICs cultured on PS-NP gels, hypothesizing that male VICs would have increased acetylation to drive expression of osteoblast-associated genes. We observed a unique trend in the males, where only larger particles affected the acetylation state (**Figure 4D-E**). Male VICs also displayed significantly higher acetylation states (**Figure 4D**) in most of our blank and PS-NP hydrogel conditions, when compared to female VICs (**Figure 4E**). Our evidence suggests that male-specific epigenetic modifications may regulate male-specific osteoblast-like differentiation on PS-NP hydrogels.

**Figure 4.**
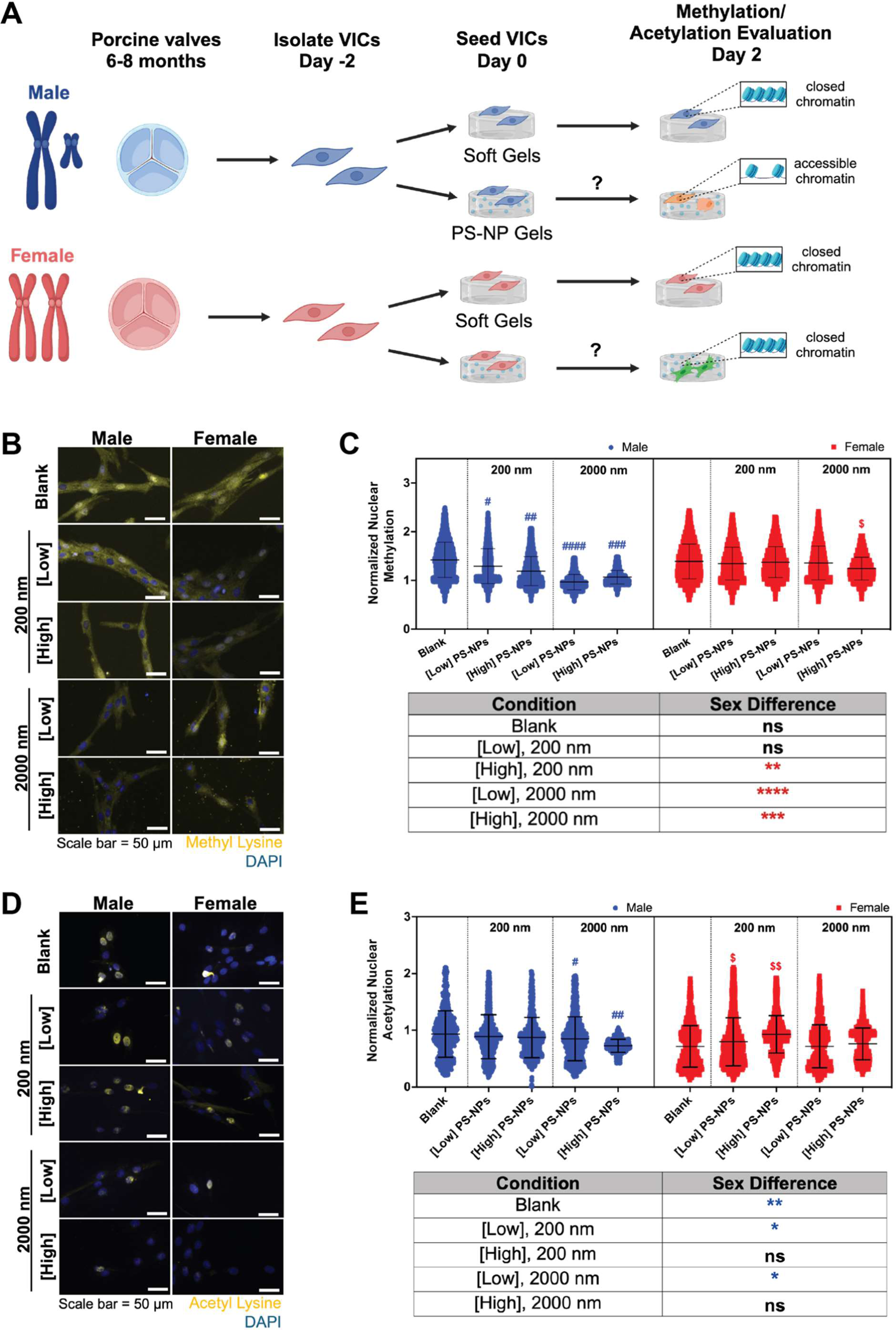
Male and female VICs exhibit sex-specific changes in global methylation and acetylation states. (**A**) Schematic describing experimental workflow for assessing sex-specific methylation and acetylation states in VICs cultured on PS-NP gels. Figure drawn with BioRender. (**B**) Representative confocal microscopy images of methylated lysine (yellow) and nuclei (blue). Scale bar = 50 μm. (**C**) Normalized methylation in male and female VICs cultured on blank hydrogels with no PS-NPs, 200 nm PS-NPs, or 2000 nm PS-NPs at low and high concentrations (0.02 mg/ml and 0.7 mg/ml, respectively). Significance determined via Cohen’s d-test (**d < 0.5, ***d<0.8, ****d<1.4 denoting significance between biological sex in the provided table, red asterisk color indicates significant increase in female groups, ^#^d < 0.2, ^##^d < 0.5, ^###^d<0.8, ^####^d<1.4 denoting significance relative to male blank with no PS-NPs, ^$^d < 0.2, ^$$^d < 0.5, ^$$$^d<0.8, ^$$$$^d<1.4 denoting significance relative to female blank with no PS-NPs). (**D**) Representative confocal microscopy images of acetylated lysine (yellow) and nuclei (blue). Scale bar = 50 μm. (**E**) Normalized acetylation in male and female VICs cultured on control hydrogels with no PS-NPs, 200 nm PS-NPs, or 2000 nm PS-NPs at low and high concentrations (0.02 mg/ml and 0.7 mg/ml, respectively). Significance determined via Cohen’s d-test (*d<0.2, **d < 0.5 denoting significance between biological sex in the provided table, blue asterisk color indicates significant increase in male groups, ^#^d < 0.2, ^##^d < 0.5, ^###^d<0.8, ^####^d<1.4 denoting significance relative to male blank with no PS-NPs, ^$^d < 0.2, ^$$^d < 0.5, ^$$$^d<0.8, ^$$$$^d<1.4 denoting significance relative to female blank with no PS-NPs).

### UTY modulates male-specific VIC phenotypes *in vivo* and *in vitro*

We next hypothesized that epigenetic modifiers coded on the Y-chromosome may contribute to male-specific VIC phenotypes *in vivo* and *in vitro*. Specifically, we posit that a Y-linked demethylase may play a key role in regulating VIC response to nanoscale cues in the extracellular matrix. We interrogated the role of the Y-linked global demethylase Ubiquitously Transcribed Tetratricopeptide Repeat Containing, Y-linked (UTY) in regulating VIC phenotype *in vivo* and variable methylation in cells cultured on PS-NP hydrogels.

We first evaluated *UTY* expression in diseased AVS valve tissue using our single cell sequencing human aortic valve datasets. In males, VIC1, VIC2, and VIC3 displayed a higher count of *UTY+* cells compared to VIC4 and VIC5 (**Supplementary Figure 8A**). Interestingly, the overall percentage of VICs expressing *UTY* decreases with disease (**Figure 5A**), subsets of VICs within each population have increased *UTY* expression levels in disease, indicating heterogeneity in *UTY* expression (**Figure 5B**). Female VICs did not display detectable *UTY* expression in both healthy and diseased samples, as expected (**Figure 5C**, **Supplementary Figure 8B**). We also characterized expression of the ubiquitously transcribed tetratricopeptide repeat, X chromosome (*UTX*) gene, revealing heterogeneous expression in both male and female VICs (**Supplementary Figure 8C-D**).

**Figure 5.**
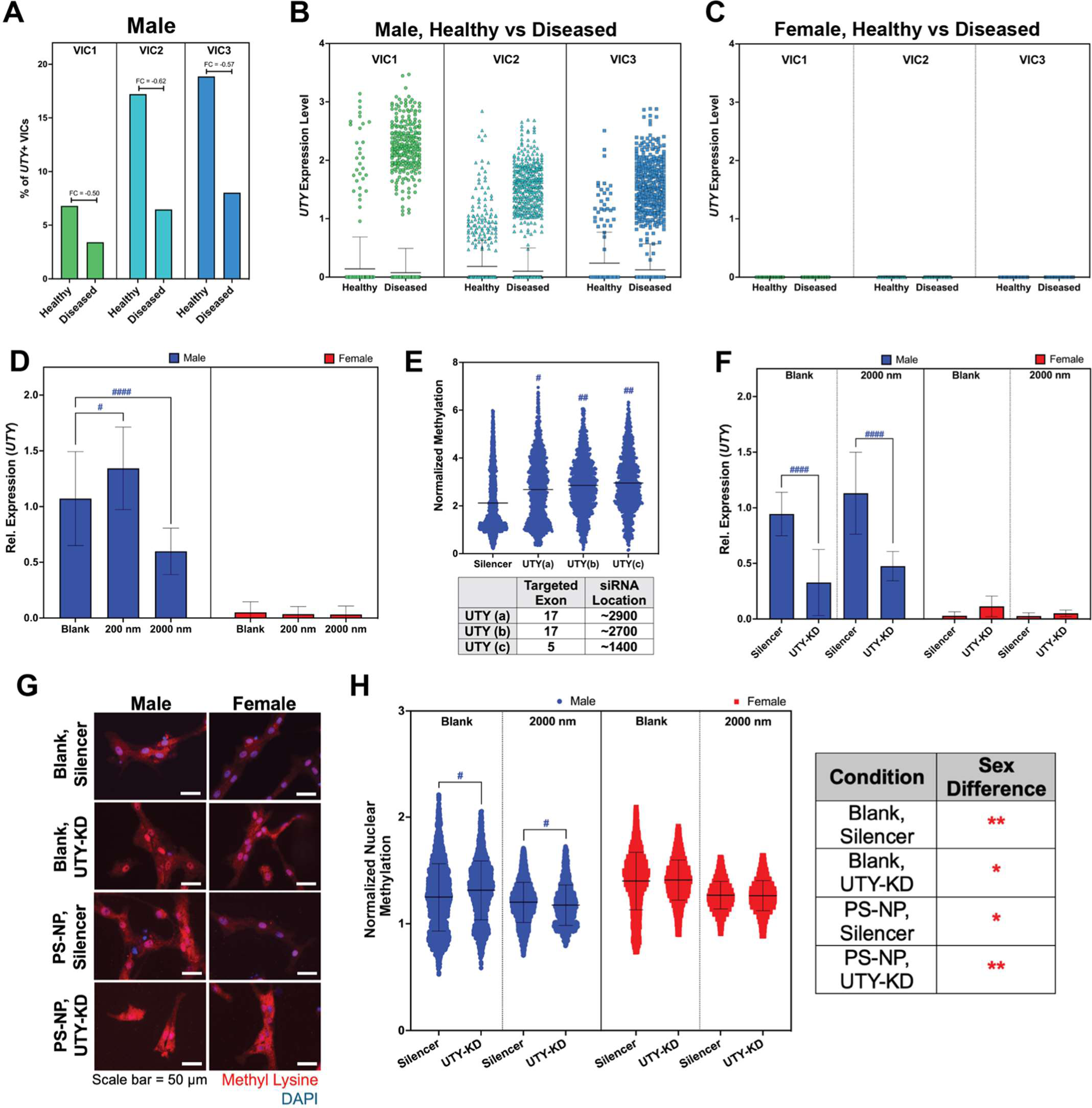
Y-linked *UTY* modulates male-specific methylation states in VICs *in vivo* and *in vitro*. (**A**) Fold change of *UTY*+ VICs in diseased human male VIC populations VIC1-3 relative to healthy male VICs. Each percentage is shown relative to the total number of VICs in each specified subpopulation. (**B-C**) *UTY* relative mRNA expression in VIC populations 1-3 in (**B**) males and (**C**) females. (**D**) Real time quantitative polymerase chain reaction (RT-qPCR) for relative expression of *UTY* in male and female VICs cultured on blank, 200 nm, and 2000 nm PS-NP hydrogels. (**E**) Normalized methylation for male VICs with three *UTY*-targeting siRNAs for exon regions 17 and 5. Significance determined via Cogen’s d-test (^#^d < 0.2, ^##^d < 0.5, ^###^d<0.8, ^####^d<1.4 denoting significance relative to male Silencer negative control siRNA vector). (**F**) RT-qPCR to validate partial silencing of *UTY* in male VICs after introduction of siRNA vector. (**G**) Representative confocal microscopy images of methylated lysine (red) and nuclei (blue). Scale bar = 50 μm. (**H**) Normalized methylation in male and female VICs cultured on blank hydrogels with no PS-NPs and 2000 nm PS-NPs. Significance determined via Cohen’s d-test (*d<0.2, **d < 0.5 denoting significance between biological sex in the provided table, blue asterisk color indicates significant increase in male groups, ^#^d < 0.2, ^##^d < 0.5, ^###^d<0.8, ^####^d<1.4 denoting significance relative to male silencer, ^$^d < 0.2, ^$$^d < 0.5, ^$$$^d<0.8, ^$$$$^d<1.4 denoting significance relative to female silencer).

To interrogate the role of *UTY* in regulating response to nanoscale stiffness cues *in vitro*, we used a small interfering RNA (siRNA) targeting the *UTY* gene to transcriptionally silence *UTY* while VICs are cultured on PS-NP hydrogels. First, we validated that *UTY* expression is male-specific on blank and PS-NP hydrogels using real time quantitative polymerase chain reaction (RT-qPCR) (**Figure 5D**). Next, we optimized our *UTY*-targeting siRNA location and targeted exon. We found that *UTY* siRNA that targets exon 5, or *UTY* (c), resulted in the largest increase in global methylation after siRNA treatment relative to the silencer negative control on a glass substrate (**Figure 5E**). For subsequent experimentation on blank and PS-NP hydrogels, we utilized *UTY* (c). After optimizing our *UTY* siRNA knockdown (*UTY*-KD) procedure, we next utilized RT-qPCR to validate that we significantly reduced relative *UTY* gene expression uniquely in male VICs cultured on PS-NP hydrogels (**Figure 5F**). Female VICs did not show significant *UTY* expression in all our conditions, further validating a male-specific reduction in *UTY* expression. As another means of validation, we used immunofluorescence to detect global methylation in male and female VIC nuclei (**Figure 5G**). As hypothesized, *UTY*-KD in male VICs led to a significant increase in methylation on blank hydrogels (**Figure 5H**). Female VICs did not increase methylation after *UTY*-KD on blank gels. On PS-NP gels, *UTY*-KD did not impact methylation in male or female VICs. Across each condition, female VICs also maintained higher levels of methylation. Together, we suggest *UTY* modulates male-specific methylation states in VICs cultured on our engineered hydrogels.

### Transcriptomic analyses unveil upregulation of pro-osteoblast pathways after *UTY* knockdown in porcine VICs on PS-NP hydrogels

Next, we hypothesized that *UTY*-KD would upregulate signaling networks associated with osteoblasts uniquely in male VICs. We performed bulk mRNA sequencing to understand the sex-specific transcriptomic responses to *UTY*-KD, given our prior observations revealing sex-specific differences in myofibroblast and osteoblast-like phenotypes, in addition to sex-specific methylation and acetylation states. In both blank and PS-NP gels, male VICs displayed a larger number of differentially expressed genes (DEGs) compared to female VICs (**Figure 6A-B**, **Supplementary Figure 9A-B**). On blank hydrogels, male and female VICs had 861 and 221 genes differentially expressed, respectively. On PS-NP hydrogels, male and female VICs had 583 and 336 genes differentially expressed, respectively.

**Figure 6.**
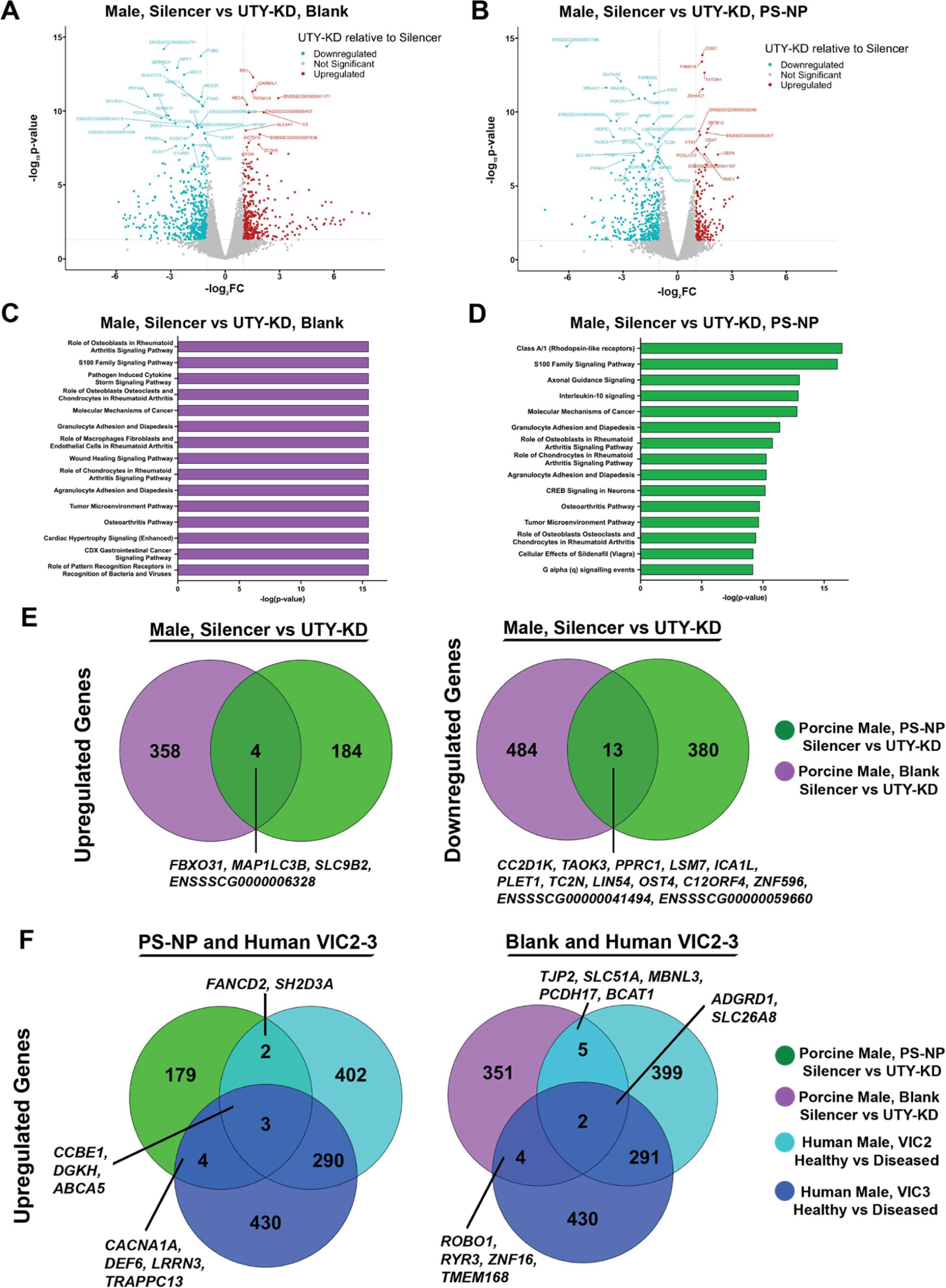
Transcriptomics analyses in porcine male VICs after *UTY*-KD unveils osteoblast-associated pathways. **(A-D)** Bulk mRNA sequencing shows differentially expressed genes in male porcine VICs (blue = downregulated, red = upregulated, gray = not significant) in (**A**) blank and (**B**) PS-NP hydrogels using DESeq2. Ingenuity pathway analysis of porcine VICs after *UTY*-KD on (**C**) blank and (**D**) PS-NP hydrogels (log_2_FC > 1, log_2_FC < -1, and p-value < 0.05). (**E**) Venn diagrams of upregulated and downregulated DEGs in porcine VICs cultured on blank and PS-NP hydrogels after *UTY*-KD. (**F**) Venn diagrams of upregulated genes in common between porcine and human transcriptomics datasets.

Ingenuity pathway analysis of male VICs with *UTY*-KD on blank (**Figure 6C**) and PS-NP gels (**Figure 6D**) showed strong associations to pathways related to osteoblasts and chondrocytes in inflammatory diseases, osteoarthritis signaling, and interleukin-4, interleukin-13, and interleukin-17 signaling, among others. Female VICs did not show any associations to pathways directly related to osteoblasts (**Supplementary Figure 9C-D).** Next, we compared the significant upregulated and downregulated DEGs in male VICs on both hydrogel substrates after *UTY*-KD (**Figure 6E**). We observed 13 downregulated genes and 4 upregulated genes in common between blank and PS-NP gels in male VICs. These data suggest *UTY* knockdown modulates genes relevant to cellular responses to microenvironmental cues.

Next, we compared our *in vitro* porcine transcriptomic dataset with our human aortic valve single-cell sequencing dataset to assess how our PS-NP hydrogel system recapitulates VIC differentiation processes *in vivo*. We compared genes upregulated after *UTY* knockdown in male porcine VICs to genes upregulated in diseased human male VICs with at least 2-fold reduction in *UTY* expression (VIC2 and VIC3). VIC1 was not included in this comparison due to its strong associations with inflammatory signaling processes, relative to VIC2 and VIC3. We observed a subset of common genes between our human VICs and porcine VICs on both blank and PS-NP hydrogels (**Figure 6F**, **Supplementary Figure 10-11**). Genes associated with pro-osteogenic differentiation and calcium homeostasis, such as *DGKH*^32,33^*, DEF6*^34,35^, and *LRRN3*^36^ were commonly identified between the porcine VICs with *UTY* knockdown and human data. We also compared upregulated genes in porcine VICs to known epigenetic modifiers^37^ and identified candidate osteogenic-associated transcriptional regulators. We observed *SUPT3H*, which has been linked to RUNX2 induced osteogenic differentiation^38–40^, in the human male VIC1-3 populations and the porcine male PS-NP condition, but not the blank hydrogel condition (**Supplementary Table 7**). Taken together, the human VIC analysis supports our PS-NP hydrogel system in the recapitulation of key aspects of stiffness-induced osteogenic differentiation via modulating *UTY* expression levels.

## Discussion

Here, we highlight that the Y-chromosome modulates male-specific VIC phenotypes to nanoscale cues in the extracellular matrix relative to female VICs (**Figure 7**). Using PEG hydrogels with polystyrene nanoparticles as a platform for cell culture, we show the protein-coding gene *UTY* may modulate male-specific demethylation in VICs during calcification in AVS. Recent studies have described the significant role of the mosaic loss of the Y-chromosome (mLOY) in male-specific cardiovascular disease progression^41,42,43^. mLOY occurs as an aging-related mutation in males, though the heterogeneity in mLOY in varying cell types and relation disease progression is not well understood. In AVS, male patients that experience mLOY have a pro-fibrotic gene expression signature in monocytes, leading to increased patient mortality after transcatheter aortic valve replacement (TAVR)^41^. Furthermore, *UTY* is also known to protect against proinflammatory phenotypes in lung tissue during pulmonary hypertension^44^. Our results provide additional evidence supporting a protective role of the Y-chromosome in disease progression, specifically valve calcification. Our results also further corroborate prior efforts implicating Y chromosome linked genes, including *UTY*, as key modulators of male-specific cell phenotypes^44,41,42^. Together, our work here provides a hydrogel cell culture platform that enables the interrogation of Y chromosome linked genes and their impact on male-specific cellular phenotypes.

**Figure 7.**
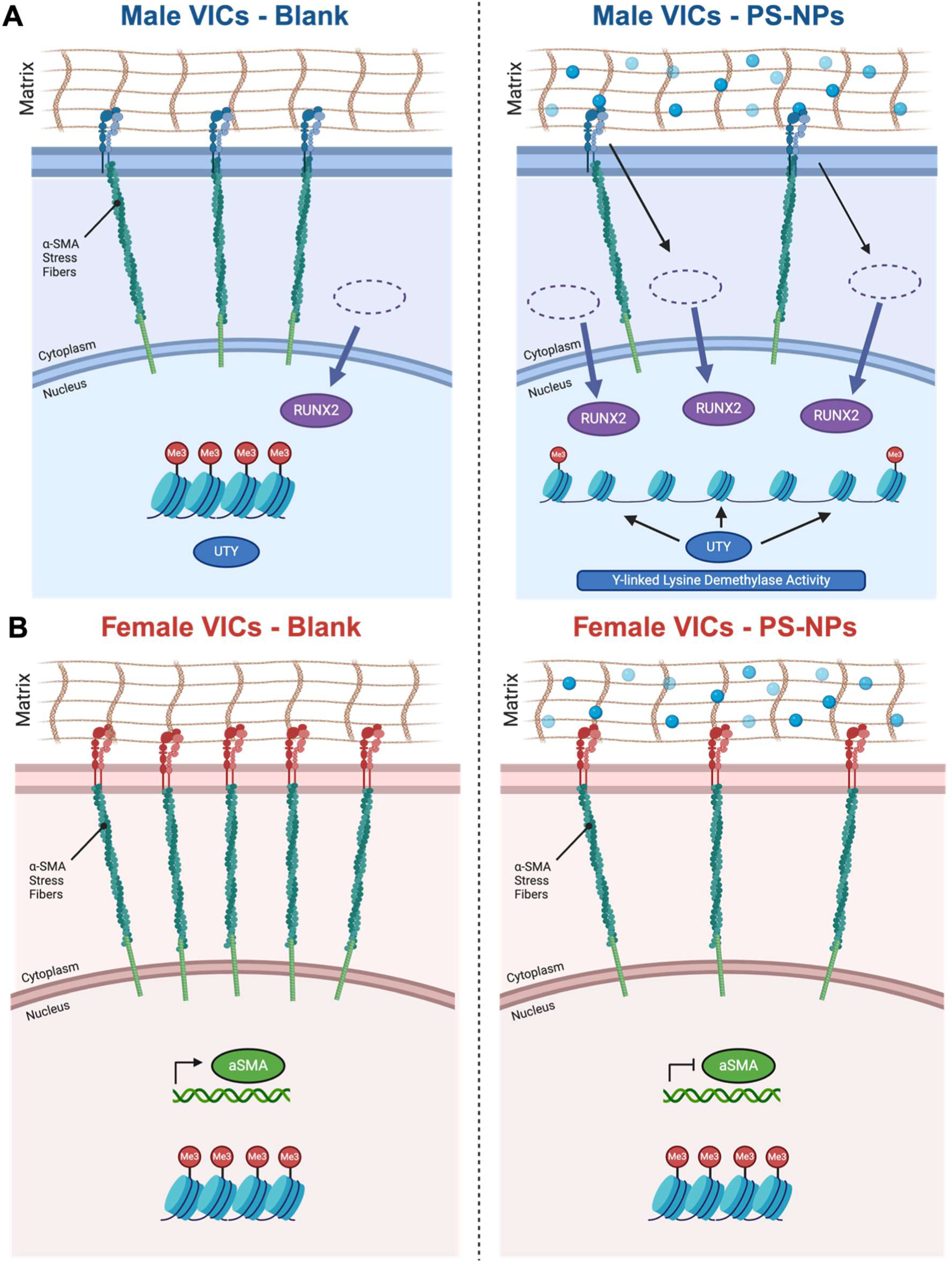
Proposed mechanism for sex-specific transcriptomic regulation in the presence and absence of nanoscale stiffness cues. At baseline conditions, male VICs maintain higher levels of Runx2 nuclear localization, while females maintain higher levels of aSMA stress fibers. (**A**) In the presence of PS-NPs, we observed increased demethylation in male porcine VICs, whereas (**B**) we only observed decreases in aSMA stress fiber presence in female porcine VICs.

To the best of our knowledge, our study is the first to directly show *UTY* influences male-specific VIC myofibroblast activation and osteoblast-like differentiation in response to extracellular matrix cues. Primary VICs from porcine sources have long been utilized to evaluate myofibroblast activation and osteogenic differentiation processes in AVS in response to different bulk substrate stiffnesses^29,45–51^. For example, in response to tissue culture plastic (elastic modulus, E ∼ 1 GPa), VICs automatically activate to myofibroblasts through the expression of ɑ-SMA stress fibers^27^. As an alternative approach, PEG hydrogel platforms have been used to study sex-specific modulation of myofibroblast activation and osteoblast differentiation states. For example, previous work has shown that VICs cultured in 2D or 3D PEG-norbornene hydrogels in the ∼1 kPa to ∼40 kPa stiffness range recapitulates sex-specific myofibroblast phenotypes observed in valve tissue^27,28^. Using a similar hydrogel system, we incorporated nanoparticles into the hydrogel matrix to mimic nanoscale features in valve tissue, as observed in our SEM images and previous work^27,28,51^. Our work suggests that RUNX2 nuclear localization in VICs responding to nanoscale features in tissue may modulate a male-specific bias toward the initiation of calcification processes.

Our study also showed *UTY* alters the VIC transcriptome on hydrogels via modulations in tri-methylation. Epigenetic modifications can occur in response to an external mechanical stimulus^18,19^, which alters the conformation of cell surface receptors and cytoskeletal network tied to the nucleus^23^. In the case of valve myofibroblasts, VICs cultured on hydrogels with varied stiffness reveal distinct chromatin structures similar to healthy and diseased valve tissue^18,19^. Building upon this work, we show that male specific methylation and acetylation states may be modulated via *UTY* demethylase activity. When evaluating methylation in males, we observed a consistent trend in decreased methylation correlating with particle size and density, indicating male myofibroblasts interact differently with nanoscale features in an extracellular microenvironment relative to female myofibroblasts. Interestingly, female VICs did not show a decrease in methylation when interacting with nanoscale cues, but did reduce methylation with larger nanoparticles, indicating other demethylases may participate in female-specific cell-particle interactions. A previous study showed that female induced pluripotent stem cells cultured with nanoparticles experienced compromised differentiation processes via disordered X chromosome inactivation, although female-specific effects of nanoparticles on pluripotent vs somatic cell cultures have yet to be compared^52^. Future efforts will need to assess the role of other demethylases in common between XY and XX cells, or potential activity of X-linked demethylases (e.g. *UTX, KDM5C*, etc.) to supplement our observations in male VICs.

Our transcriptomics analyses revealed UTY as a modulator of osteogenic associated genes relevant to calcification. We found reduced *UTY* expression *in vitro* led to the upregulation of numerous pathways broadly associated with osteoblast signaling, inflammation, and wound healing, which mirror upregulated pathways in human male VIC2-3 populations with reduced *UTY* expression. Of note, we found epigenetic modifiers *SUPT3H*, *DEF6, and LRRN3* were uniquely upregulated in the porcine PS-NP *UTY*-KD condition along with human male VIC2-3. Previous work has shown *SUPT3H* promotes H3 acetylation and subsequent Runx2 transcription^38–40^, promoting osteoblast differentiation. Prior studies have also linked demethylation to accelerated osteogenic differentiation processes via *DEF6*^34,35^ and *LRRN3*^36^. Our analysis also identified upregulation of matrix remodeling and calcification-associated genes in VICs with *UTY* knockdown, including *CCBE1*^53^ and *DGKH*^32,33^. We also identified several pro-fibrotic genes and genes involved in myofibroblast activation and age-related cardiovascular diseases, including *MBNL3*^54–52^ and *ROBO1*^53^. Collectively, we posit *UTY* may promote male-specific calcification processes in the human aortic valve via multiple gene signaling networks. Our work also provides a steppingstone to future validation studies to further understand how sex chromosomes and the extracellular microenvironment synergistically contribute to somatic cell phenotypes in health and disease.

There are several limitations of the PS-NP hydrogel platform described in our study. First, our study did not fully decouple the effects of sex hormone imprinting from sex chromosomes^55–57^, as only our male pigs were gonadectomized at birth. We envision future work where VICs from gonadectomized males and female animal models can be used to further study the effects of the sex chromosome on VIC phenotype. Secondly, mechanical and biochemical cues synergistically modulate the VIC to myofibroblast to osteoblast-like cell transition during AVS. Future cell culture platforms can be adapted to include inflammatory factors secreted from macrophages to evaluate epigenetic modifications in response to inflammatory cytokines and nanoscale stiffness cues. Third, our study also evaluated cell phenotypes after two days in culture. Acknowledging that AVS is a time-dependent progressive disease, we suggest future studies where longer timepoints are used to evaluate sex-specific chromatin structures (e.g. after days to weeks in culture) as VICs acquire a more permanent myofibroblast phenotype reflective of diseased valve tissue. Fourth, future efforts to use spherical calcium phosphate particles instead of PS-NPs may be used to further enhance the physiological relevance of the *in vitro* culture platform and evaluate effects of *UTY* and other Y-linked genes on calcification^58^.

Taken together, our study implicates the importance of sex chromosome-linked genes in the progression of disease-driving phenotypes in AVS. Our bioinspired hydrogel platform provides evidence that nanoscale cues in the extracellular matrix influence sex-specific phenotypes in VICs, and broadly suggests biomaterials are useful tools to answer research questions related to sex-specific biology^59,60^. We posit that Y-linked epigenetic regulators, such as *UTY*, regulate male-specific osteoblast-like phenotypes, contributing to earlier onset of valvular calcification in males. Future studies will explore the interplay between mechanical stimuli and sex-specific epigenetic regulation in males and females.

## Materials and Methods

### Nanoparticle hydrogel fabrication

8-arm, 40 kDa polyethylene(glycol)-norbornene (PEG-Nb) was synthesized as previously described^61^. Hydrogel precursor solutions were made mixing together 4% (w/v) PEG-Nb, 5 kDa PEG-dithiol crosslinker (2.96 mM, JenKem), CRGDS peptide (2 mM, Bachem), phosphate buffered saline (PBS, Gibco, Cat. No. 14-190-250), and photoinitiator lithium phenyl-2,4,6-trimethyl-benzoylphosphinate (1.7 mM, Sigma, Cat. No. 900889). To incorporate nanoparticles into hydrogel precursor solutions, polystyrene nanoparticle solutions, we utilized 1% wt/vol aqueous solutions of 200 nm (Nanocs, Cat. No. PS01-200) or 2000 nm (Nanocs, Cat. No. PS01-2u) polystyrene nanoparticles (PS-NPs) instead of PBS in the hydrogel precursor formulation. All hydrogels were made utilizing a 0.99:1 thiol-to-ene ratio. Vapor deposition was used to functionalize 12 mm glass coverslips (Mercedes Scientific, Cat. No. MER0012) and 25 mm glass coverslips (Mercedes Scientific, Cat. No. MER0025) with thiol groups to allow for hydrogel adhesion. Briefly, coverslips were placed in an autoclave jar along with 100 µL of mercaptopropyltrimethoxysilane (MPTS, Sigma-Aldrich, Cat. No. 175617) in a 60°C oven overnight for thiol functionalization. Glass slides were coated with Sigmacote (Sigma-Aldrich, Cat. No. SL2) to create a hydrophobic glass surface during hydrogel fabrication. Either 11.3 µL or 65 µL of hydrogel precursor solution was pipetted onto the Sigmacote glass slide and covered with a 12 mm or 25 mm coverslip, respectively. The hydrogels were photopolymerized at 4 mW/cm^2^, removed with a razor blade, sterilized with 5% (v/v) isopropyl alcohol/PBS solution, and rinsed three times with PBS. After the final PBS rinse, cell culture media containing Media 199 (Gibco, Cat. No. 11043023) supplemented with 1% Fetal Bovine Serum (Gibco, Cat. No. 16000069), and 50 units penicillin, and 0.05 mg/ml streptomycin (Sigma Aldrich, Cat. No. P4458) was added to the hydrogels for overnight swelling prior to cell culture.

### Rheology

Hydrogels containing nanoparticles were prepared as described above for rheological measurements. Storage (G’) and loss (G’’) moduli were measured using a DHR-3 rheometer (TA Instruments) with an 8 mm parallel plate geometry. Oscillatory shear rheology was used with an amplitude of 1% and frequency of 1 Hz^26^. To obtain the elastic modulus (E), the following formula was used: E = 2 * G’ (1 + v) where v is Poisson’s ratio. We utilized a value of v = 0.5, assuming G’ >>> G’’ for viscoelastic hydrogels.

### Human Aortic Valve Tissue Digestion and Tissue Preservation

Human aortic valve tissue samples were obtained from the National Disease Research Interchange (NDRI) with UC San Diego Institutional Review Board approval (UCSD IRB 804209). Aortic valves were excised from human hearts (post-mortem interval < 48 hours) and washed with Earle’s Balanced Salt Solution (EBSS, Gibco, Cat. No. E2888). Valve leaflets without evidence of calcification were categorized as “Healthy.” We also categorized mid and late stage AVS patient samples as “Diseased” to better represent the heterogeneity of disease presentation in patients (**Supplemental Table 1**). Briefly, valve leaflets were then digested with Liberase TM (Roche, Cat. No. 05401119001) for 30 minutes before quenching with media for centrifugation. The resulting cell pellet was treated with Red Blood Cell Lysis Buffer (Invitrogen, Cat. No. 00433357) for 1 minute before quenching with media and centrifugation. Cells were counted manually with a hemocytometer before subsequent preparation for single cell sequencing. Valve leaflets were also placed in 4% paraformaldehyde and stored at 4C for paraffin embedding, sectioning, and subsequent scanning electron microscopy.

### Scanning Electron Microscopy (SEM) on Hydrogels and Human Valve Tissue and quantification of particle size distribution

For scanning electron microscopy (SEM) of hydrogels, samples were prepared on coverslips, frozen in liquid nitrogen, and lyophilized. SEM was performed using a JSM-6010LA instrument (JEOL Ltd, CU Anschutz) using manufacturer settings. For SEM on human valve tissue, samples were fixed, paraffin embedded, and sectioned onto glass slides. Samples were sputter coated with iridium for 30 seconds at 75% power and imaged using a FEI Quanta FEG 250 on manufacturer settings. Average calcium particle size was determined from two biological replicates for each sex in the disease condition using ImageJ as shown in **Supplementary Figure 6A**.

### Atomic Force Microscopy (AFM)

For atomic force microscopy, hydrogels mounted on coverslips were immersed in PBS and analyzed using a JPK NanoWizard 4a instrument (JPK Instruments). Hydrogels were probed using CB3 cantilevers with triangular tips (Nanosensors, qp-BioAC-CI) until an initial force of 1 nN was reached. Calibration of the cantilever was made applying the thermal noise method. Force spectroscopy was performed at each node of a square grid distributed over a 20 µm by 20 µm grid. Force–displacement (F–d) data were collected using a peak load of 5 nN at a displacement rate of 10 µm/s. Force–deformation (F–δ) data were calculated using F–d curves by subtracting the initial cantilever deflection. To determine Young’s modulus E at each node, the loading portion of each F–δ curve was fit to an analytical model for a rigid conical tip in contact with an elastic half-space (38), i.e., F=(2/π)(E/1−ν^2^)(tanα)δ^2^, using angle α determined from SEM images, an assumed Poisson’s ratio for the hydrogel (ν = 0.5), and E as the sole fitting parameter.

### Porcine valvular interstitial cell isolation and culture

Aortic valves (AVs) were dissected from 5-8-month-old male or female pigs (Midwest Research Swine) to obtain valvular interstitial cells (VICs). For each isolation, four to eight porcine hearts were separated by sex and pooled. Each batch of VICs obtained from an isolation is considered as a biological replicate, in which 2-3 biological replicates were used for **Figures 2-6**. After dissection, AV leaflets were submerged and rinsed in EBSS containing 50 units penicillin and 0.05 mg/ml streptomycin. Next, leaflets were digested in EBSS containing 250 units of Collagenase type 2 (Worthington Biochemical, Cat. No. LS004176) for 30 minutes at 37°C under constant agitation at 100 rpm. After, collagenase solution was aspirated from the AV leaflets to remove endothelial cells. A secondary digestion with fresh collagenase solution was performed for 60 minutes under agitation at 100 rpm. To collect VICs, the remaining tissue was scraped against a 100 µm cell strainer, washed with VIC media, then centrifuged at 400 g for 10 minutes. The supernatant was aspirated and replaced with fresh VIC expansion media, comprised of Media 199 supplemented with 15% FBS, and 50 units penicillin and 0.05 mg/ml streptomycin. Freshly isolated VICs were cultured in VIC expansion media with media changes every other day until confluency. VICs were frozen down at passage 1. All experiments utilized passage 2 VICs at 35,000 cells/cm^2^. To reduce proliferation during experimentation, passage 2 VICs were cultured in VIC media containing 1% FBS.

### Small-interfering RNA (siRNA) Gene Knockdown

Gene knockdowns in the porcine VICs were conducted using small interfering RNAs (siRNAs). We utilized Ubiquitously Transcribed Tetratricopeptide Repeats, Y-linked (*UTY*) siRNA (Ambion, Cat. No. 4392421, ID: s14740). On Day 0, cells were seeded at 42,000 cells/cm^2^ coverslips in 1% VIC media, without antibiotics or antifungal supplements, and cultured overnight. We increased cell seeding density per manufacturer recommendations for siRNA experiments. On Day 1, *UTY* siRNAs and Silencer negative control (Ambion, Cat. No. AM4635) were incubated with Lipofectamine 3000 Transfection Reagent (Thermo Scientific, Cat. No. L3000008) for 15 minutes at room temperature before adding it to fresh 1% VIC media without antibiotics. The final siRNA concentration for the 12 mm and 25 mm hydrogels was 20 pmol and 100 pmol, respectively. On Day 2, the samples were monitored without media change. On Day 3, the samples were fixed for immunofluorescence and lysed for RNA collection. In total, *UTY* siRNA treatment lasted 48 hours.

### Immunofluorescence, Imaging, and Analysis

Cells were cultured on NP hydrogels for 48 hours before fixing with 4% (w/v) paraformaldehyde (Electron Microscopy Sciences, Cat. No. 15710) for 20 minutes. Afterwards, cells were permeabilized with 0.1% (w/v) Triton-X-100 (Sigma Aldrich, Cat. No. 93443) for one hour. Permeabilization buffer was removed and added 5% (w/v) bovine serum albumin (BSA, Sigma Aldrich, Cat. No. A8327) as blocking buffer overnight at 4°C. The following primary antibodies were diluted in blocking buffer and incubated for one hour at room temperature: α-smooth muscle actin (aSMA, 1:300, Abcam, Cat. No. ab7817), runt-related transcription factor 2 (RUNX2, 1:300, Abcam, Cat. No. ab23981), Methyl-Lysine (MeK, 1:350, Novus Biologicals, Cat. No. NB600824), and acetylated lysine (AcK, 1:350, Abcam, Cat. No. ab190479). Samples were rinsed with 0.05% (v/v) Tween-20 (Sigma Aldrich, Cat. No. P1379) for 5 minutes. The following secondary antibodies were diluted in blocking buffer and incubated for one hour at room temperature protected from light: AlexaFluor 488 goat anti-mouse (1:200, Thermo Scientific, Cat. No. A11001), AlexaFluor 647 goat anti-rabbit (1:200, Thermo Scientific, Cat. No. A21245), or AlexaFluor 647 donkey anti-rabbit (1:200, Thermo Scientific, Cat. No. A32790). Nuclei were stained with DAPI (1:500, Roche, Cat. No. 10236276001) and cytoplasm were stained with CellMask Orange (1:5000, Fisher Scientific, Cat. No. H32713). Sample coverslips were rinsed with PBS and transferred to a glass-bottom well plate (CellVis, Cat. No. P24-1.5H-N) for imaging on a Nikon Eclipse Ti2-E. Fluorescence was quantified after modifying a freely available MATLAB code^27^, and fluorescence channel exposure was adjusted consistently in all conditions for representative image clarity. Alpha smooth muscle actin intensity ratios were calculated by normalizing α-SMA intensity with CellMask cytoplasmic intensity. Nuclear RUNX2 intensity ratios were calculated by normalizing RUNX2 nuclear intensity to cytoplasmic intensity. Nuclear methylated lysine ratios were calculated by normalizing MeK nuclear intensity to cytoplasmic intensity. Nuclear acetylated lysine ratios were calculated by normalizing AcK nuclear intensity to DAPI intensity.

### RNA Isolation and RT-qPCR

RNA was collected at specified timepoints using the RNeasy Micro Kit (Qiagen, 74004) following the manufacturer’s protocol. Gels on 25 mm coverslips were inverted onto lysis buffer for 3 minutes then rinsed with 70% (v/v) ethanol to collect lysed cellular material. After following manufacturer’s protocol for RNA isolation, samples were analyzed for mRNA concentration using a NanoDrop 2000 spectrophotometer (Thermo Scientific). Sample concentrations were normalized to 1 ng/µL during cDNA preparation using the iScript Synthesis Kit (Bio-Rad, Cat. No. 1708841), following manufacturer’s protocol. To measure relative gene expression, iQ SYBR Green Supermix (Bio-Rad, Cat. No. 1708882) was mixed with primer sequences found in Supplemental Table 2. Cq values were determined using a CFX384 iCycler (Bio-Rad). All gene expression calculations were normalized to *RPL30* expression.

### Bulk RNA Sequencing and Analysis (Porcine)

Prior to bulk paired-end RNA sequencing, RNA was isolated as described above. RNA quality was assessed by Agilent TapeStation 4200 and RNA quantity was determined by QuBit 3.0 Fluorometer. Approximately 200 ng of RNA per sample was used for sequencing. The Sanford Consortium for Regenerative Medicine’s Genomics Core conducted cDNA library preparation and indexing using the Illumina Stranded mRNA kit and IDT Illumina RNA UD Indexes, respectively. Sequences were checked for quality control using fastqc (v0.12.1) and subsequently removed adapter sequences using trimmomatic (v0.39) if adapter sequence content exceeded the threshold identified by fastqc. Alignment to the porcine genome (Sscrofa11.1) was conducted using HISAT2 (v2.2.1). Sequencing run statistics and genomic mapping accuracy are shown in **Supplementary Table 3**. Subsequent differential gene expression analysis was conducted in R (4.3.2) and R Studio (2023.12.1) using DESeq2 (Release 3.19) before we filtered for genes with a false discovery rate above 0.1. Upregulated DEGs were identified by log_2_FoldChange>1, p-value<0.05, and false discovery rate<0.1. Downregulated DEGs were identified by log_2_FoldChange<-1, p-value<0.05, and false discovery rate<0.1. **Supplementary Figure 12** shows a heatmap of the most highly differentially expressed genes across each comparison. We utilized gene ontology analysis features in Ingenuity Pathway Analysis (Qiagen) for gene enrichment analyses of upregulated and downregulated genes in the porcine and human sequencing samples.

### Single Cell Sequencing, Integration Analysis, and Statistical Analysis (Human)

We conducted a sex-separated re-analysis of previously published single cell sequencing data for healthy and calcific aortic valve disease (CAVD) patients^62^. We supplemented the analysis with two of our human valve sample patients diagnosed with aortic valve stenosis to increase biological replicates for each condition. Briefly, we prepared our human aortic valve sample library using Chromium Next GEM Single Cell 3’ Library Construction V3 Kit, following the Single Cell 3′ v3.1 Reagent Kits User Guide (10X Genomics, Document CG000204). We targeted 2000-5000 cells and sequenced at approximately 5000 reads per cell. The libraries were sequenced on the NovaSeq X Plus platform. Data were processed with Cell Ranger (v7.0.2) and raw counts were used to map the reads to the human reference genome (hg19). The outputted gene expression matrix was analyzed using the Seurat pipeline (v5.1.0)^63–67^. We excluded cells with less than 200 unique genes or more than 5 percent mitochondrial transcripts. Overall, we analyzed 40,352 cells across 6 patients (**Supplementary Table 4-6**).

We next applied Seurat’s NormalizeData function on the raw counts matrix; the function divides the feature counts of each cell by the total counts for that cell, scales that value by 10^6^, and applies a natural-log transformation. Next, we implemented the FindVariableFeatures function to choose the top 2000 highly variable genes using the “vst” selection method. After mean centering and scaling we carried out Principal Component Analysis (PCA) on a merged gene expression matrix of all 40,352 cells. We used an elbow plot analysis to select 30 principal components as the cutoff dimension of our data set. To batch correct for patient samples collected from different studies, we performed the Seurat scRNA-seq integration to match shared cell types across different datasets. To identify different cell-types, we performed unsupervised clustering via the Lovain algorithm on k-nearest neighbors with the FindNeighbors and FindClusters (parameters k=30 and resolution=0.4) commands in Seurat. The functions returned 8 cell clusters which were visualized on a Uniform Manifold Approximation and Projection (UMAP). To label individual cell clusters, we performed a Wilcox test using the FindAllMarkers function in Seurat to plot the top 10 most differentially expressed genes (p value < 0.001 and log2 fold change > 1) for each cluster (**Supplementary Figure 3**). Additionally, we used the top 10 conserved markers (minimum p value < 0.001 and log2 fold change > 0.5) across healthy and diseased samples to accurately label cell-types (**Supplementary Figure 2**). Gene expression was visualized using FeaturePlot, DoHeatMap, and VlnPlot functions in Seurat.

### Statistical Analysis

For immunofluorescence studies, one-way ANOVA with Tukey’s posttests were conducted. We set a threshold of significance cutoff at P < 0.0001. Due to the experimental sample size (N > 1000 cells), we sought to reduce statistical bias by conducting a Cohen’s d-value test to determine significance independent of sample size. Thresholds of significance were represented by the following key: *** = d > 0.8, ** = d > 0.5, * = d > 0.2 for between sex significance; ### = d > 0.8, ## = d > 0.5, # = d > 0.2 for male group significance; and $$$ = d > 0.8, $$ = d > 0.5, $ = d > 0.2 for female group significance. For RT-qPCR studies, one-way ANOVA with Tukey’s posttests were conducted. For **Figure 2**, data is shown as the mean ± standard deviation, as calculated in GraphPad PRISM.

## Acknowledgments

We would like to thank the Stem Cell Genomics Core at the Sanford Stem Cell Institute for providing sequencing services. This publication includes data generated at the UC San Diego IGM Genomics Center utilizing an Illumina X Plus that was purchased with funding from a National Institutes of Health SIG grant (#S10 OD026929). We also acknowledge the help and resources of Dr. Guillaume Castillon at the Electron Microscopy Core. This work was performed in part at the San Diego Nanotechnology Infrastructure (SDNI) of UCSD, a member of the National Nanotechnology Coordinated Infrastructure (NNCI), which is supported by the National Science Foundation (Grant ECCS-2025752). Microscopy was performed at the Nikon Imaging Center at UC San Diego. We would like to thank Dr. Peng Guo and Dr. Richard Sanchez for their support on microscopy experiments.

## Funding

National Institutes of Health, National Heart, Lung, and Blood Institute (NHLBI) grant R00 HL148542 (RMG)

San Diego ARCS Foundation (RMG)

National Institutes of Health T32 Fellowship/San Diego Match Fellowship (TB)

National Institutes of Health, NHLBI grant R00 HL148543 (NEFV)

National Science Foundation Graduate Research Fellowship Program (NEFV)

National Institutes of Health, NHLBI R01 HL171197 (KSA)

National Institutes of Health, NHLBI R01 HL147064 (LM)

National Institutes of Health, NHLBI K25 HL148386 (BP)

National Institutes of Health, NHLBI R01 HL169578 (BP)

National Institutes of Health, NHLBI R21 AG080257 (BP)

Ludeman Family Center for Women’s Health Research seed grant (BP)

American Heart Association 23CDA1052411 (BP)

John Patrick Albright (LM, BP)

National Institutes of Health, NHLBI R00 HL148542 (BAA)

National Institutes of Health DP2 HL173948 (BAA)

Chan Zuckerberg Initiative Science Diversity Leadership Award (BAA)

American Heart Association 23CDA942253 (BAA)

## Supplementary Materials

**Supplementary Figure 1:**
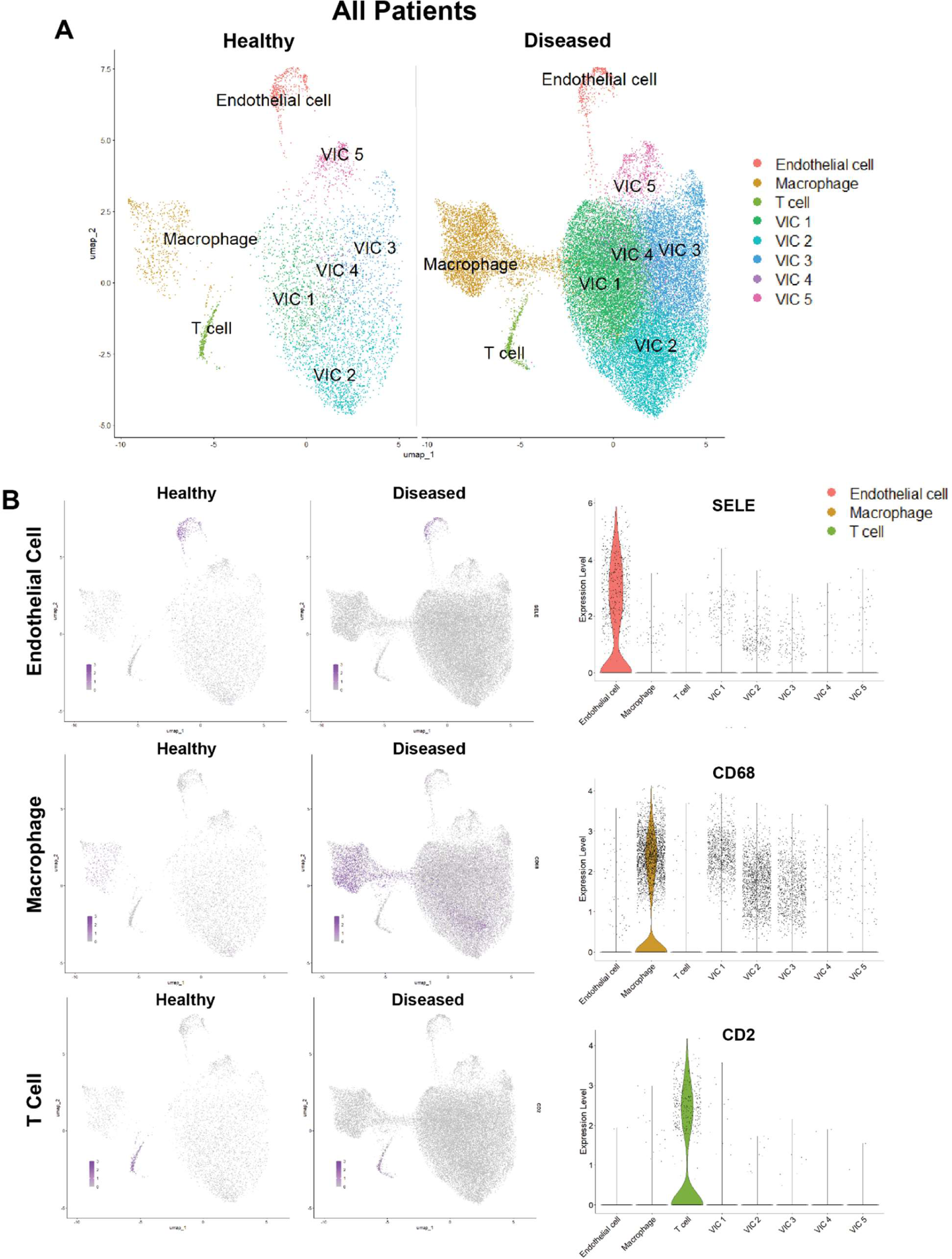
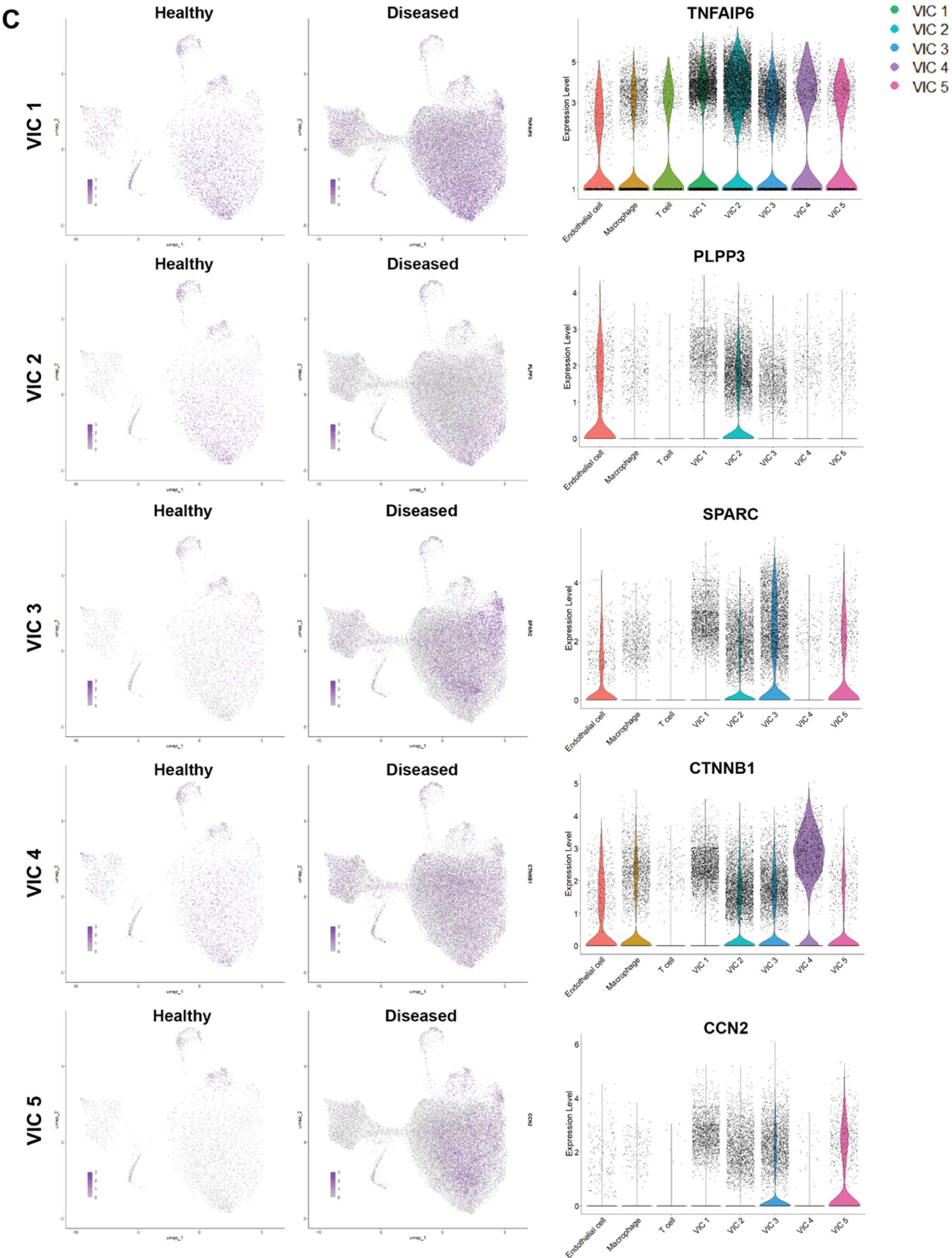
Human aortic valve cell type distribution and heterogeneity across sex and disease. (**A**) UMAP clustering and cell population distribution in healthy and diseased patient samples for males and females. Biomarkers used to identify distinct clusters in (**B**) endothelial cell and immune cell populations and (**C**) VIC populations.

**Supplementary Figure 2:**
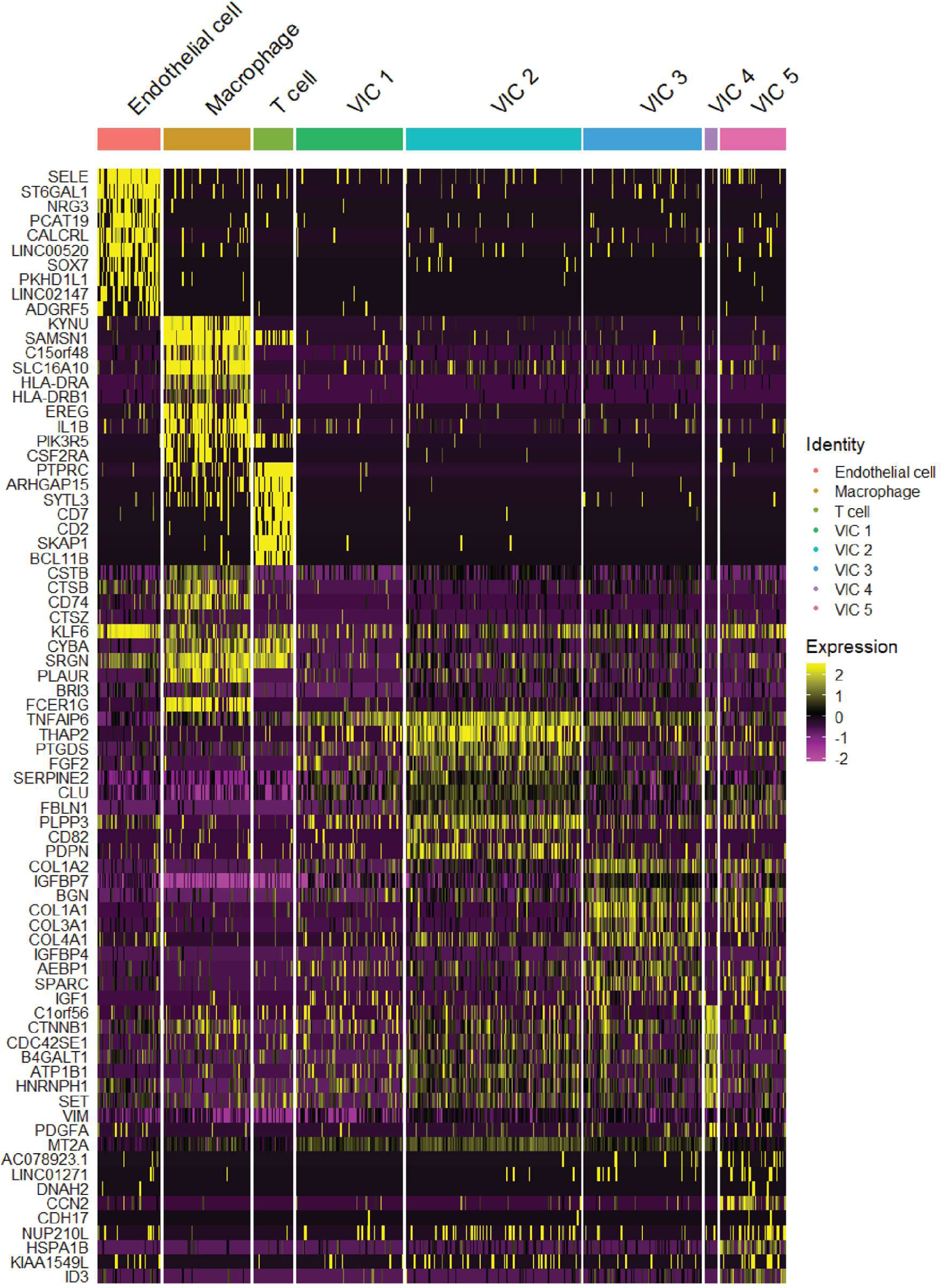
Heatmap of conserved biomarkers across all cell clusters for all patients.

**Supplementary Figure 3:**
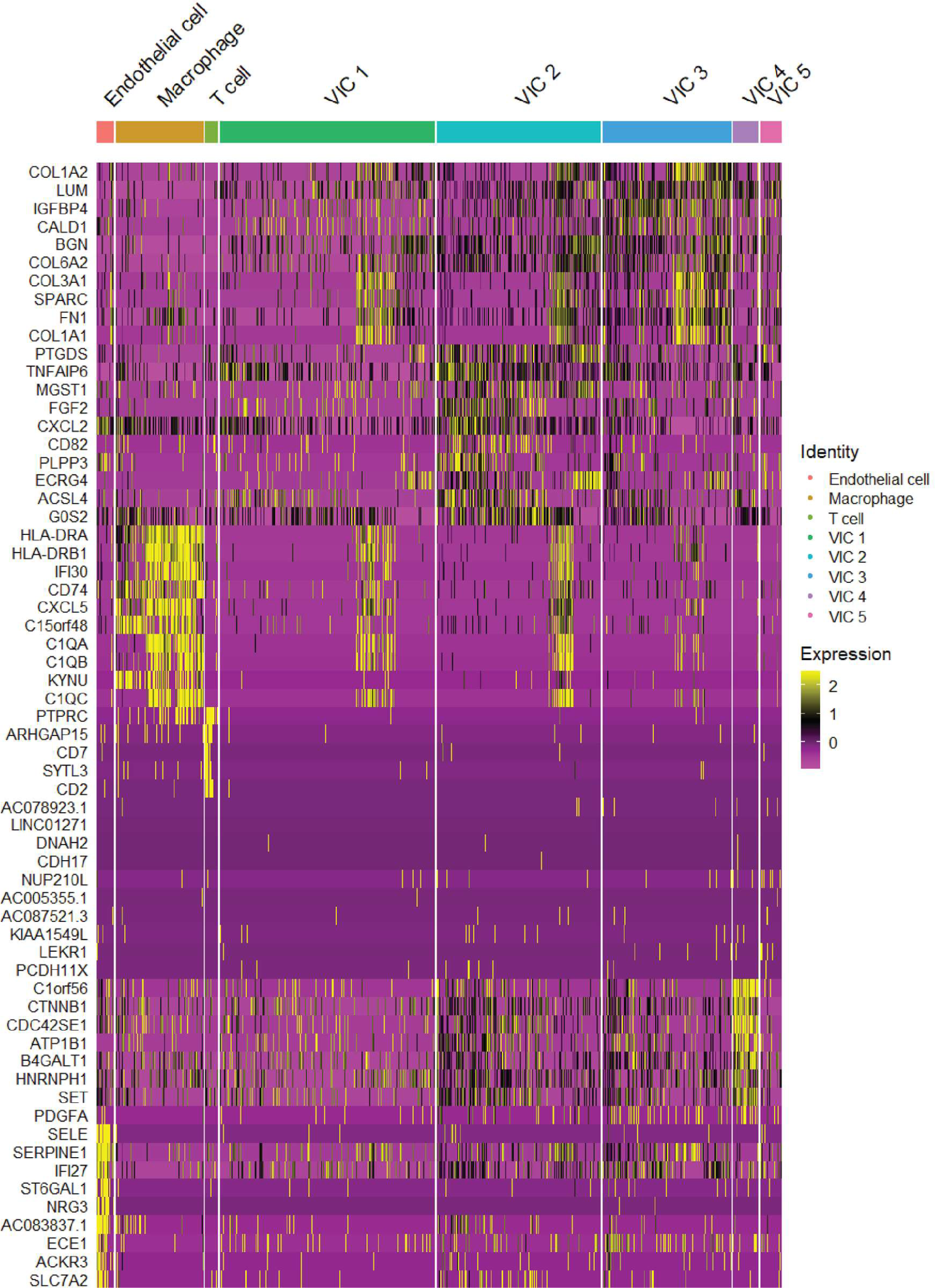
Heatmap of distinct biomarkers across all cell clusters for all patients.

**Supplementary Figure 4:**
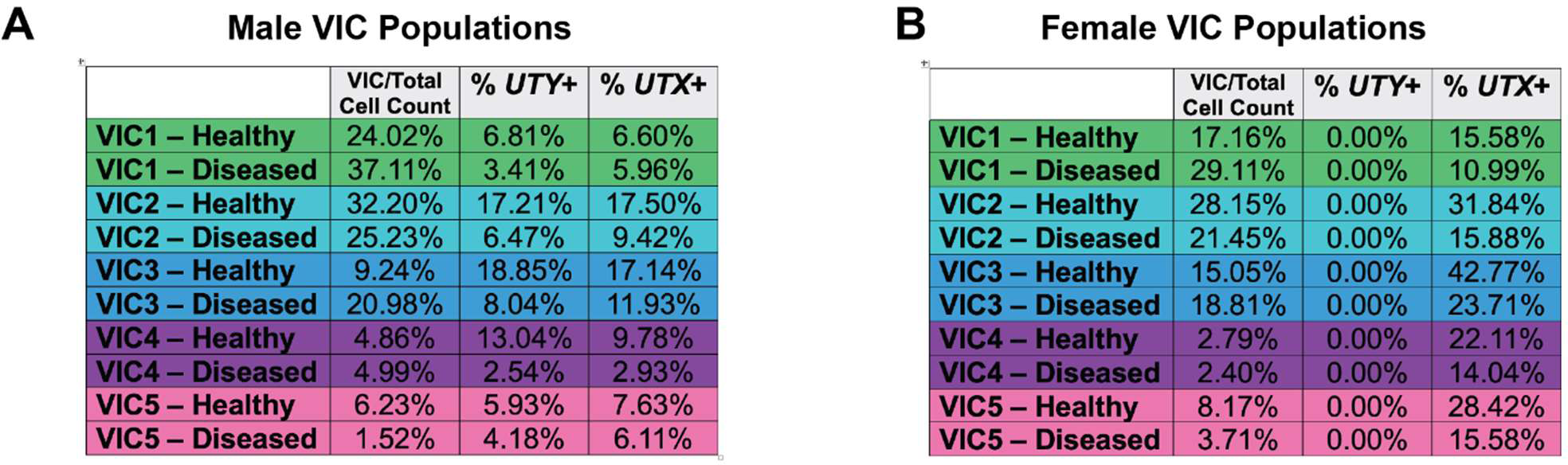
Distribution of VIC populations in (A) male and (B) female patients, in healthy and diseased states.

**Supplementary Figure 5:**
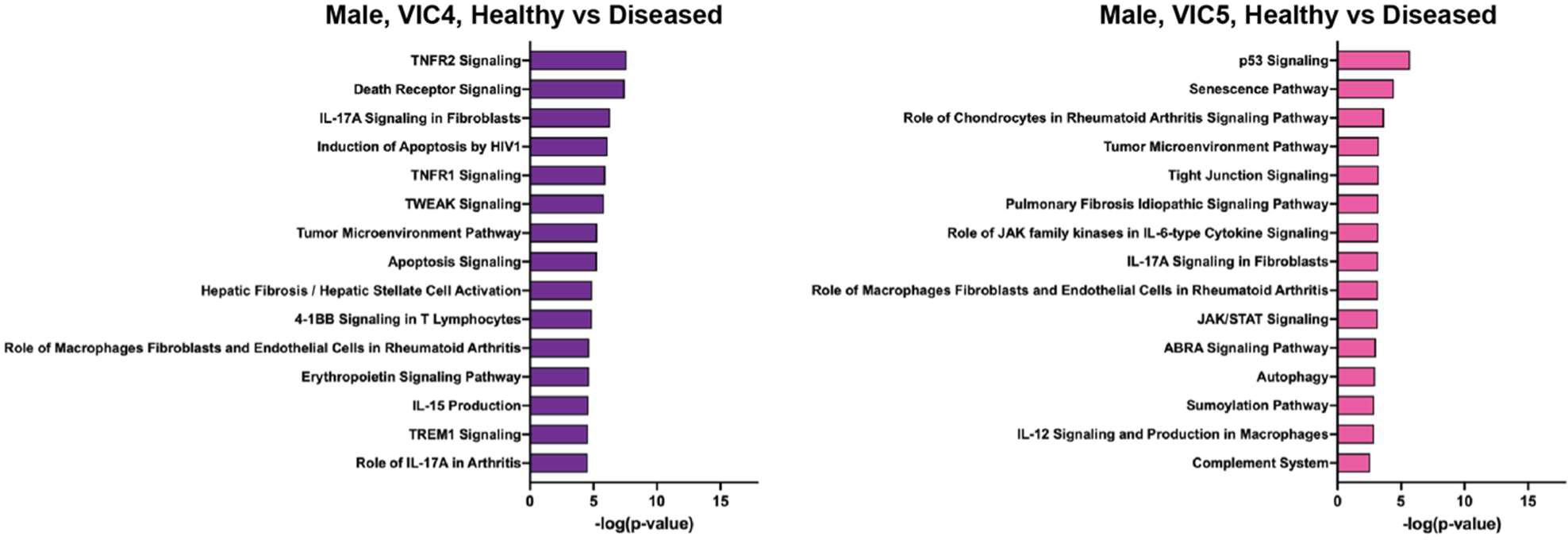
Ingenuity pathway analysis of male VIC4 and VIC5 populations.

**Supplementary Figure 6:**
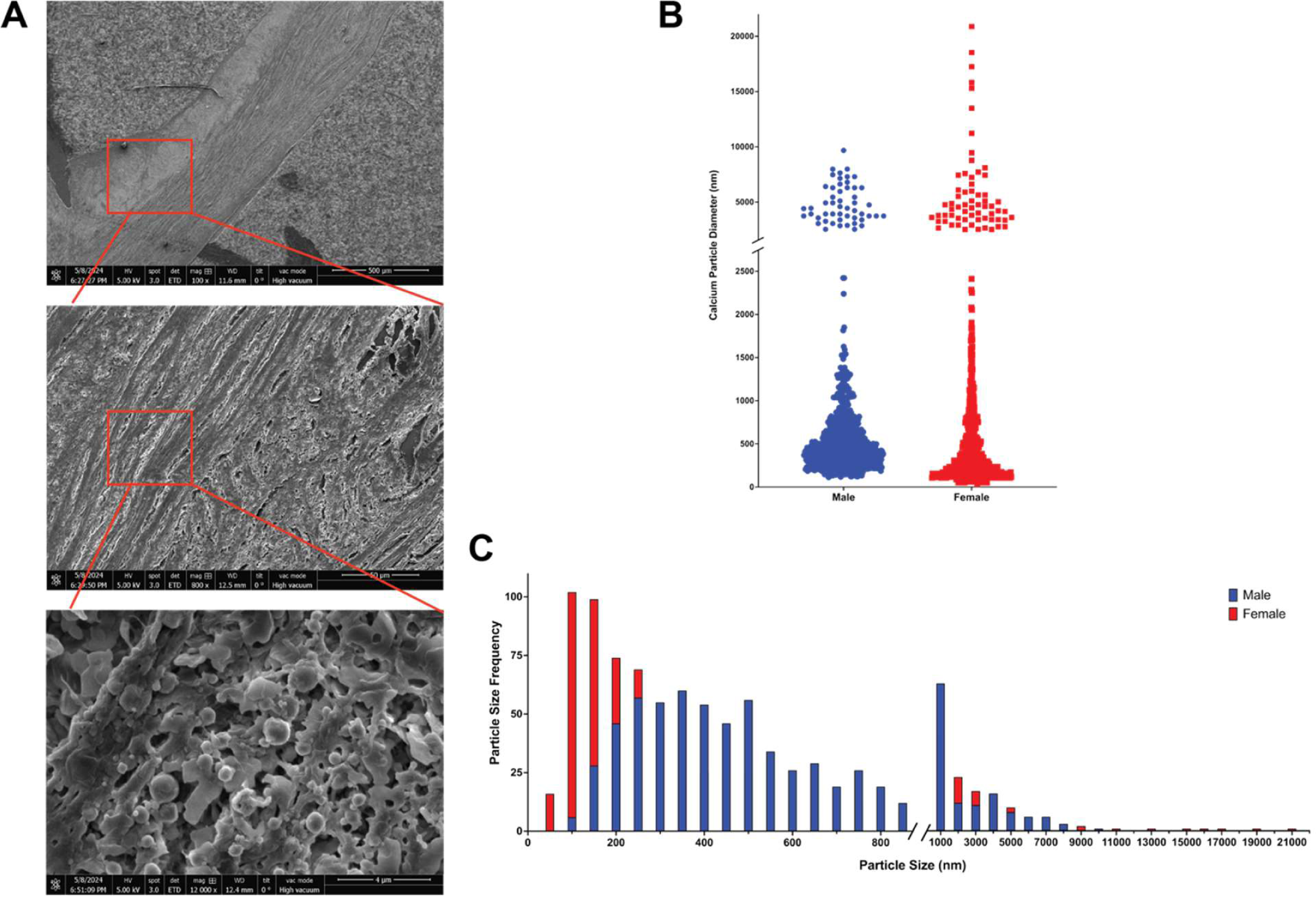
Quantification of particle size and distribution in diseased human aortic valve tissue. (**A**) SEM images of representative male aortic valve tissues to localize calcium phosphate particles. (**B**) Calcium particle diameter in diseased, age-matched male aortic valve tissue relative to female aortic valve tissue (n=2 male, n=2 female, n>600 particles, ****p < 0.0001). Diseased patients include mid and late stage AVS. (**C**) Particle diameter frequency distribution in male and female valve tissues (n=2 male, n=2 female, n>600 particles).

**Supplementary Figure 7:**
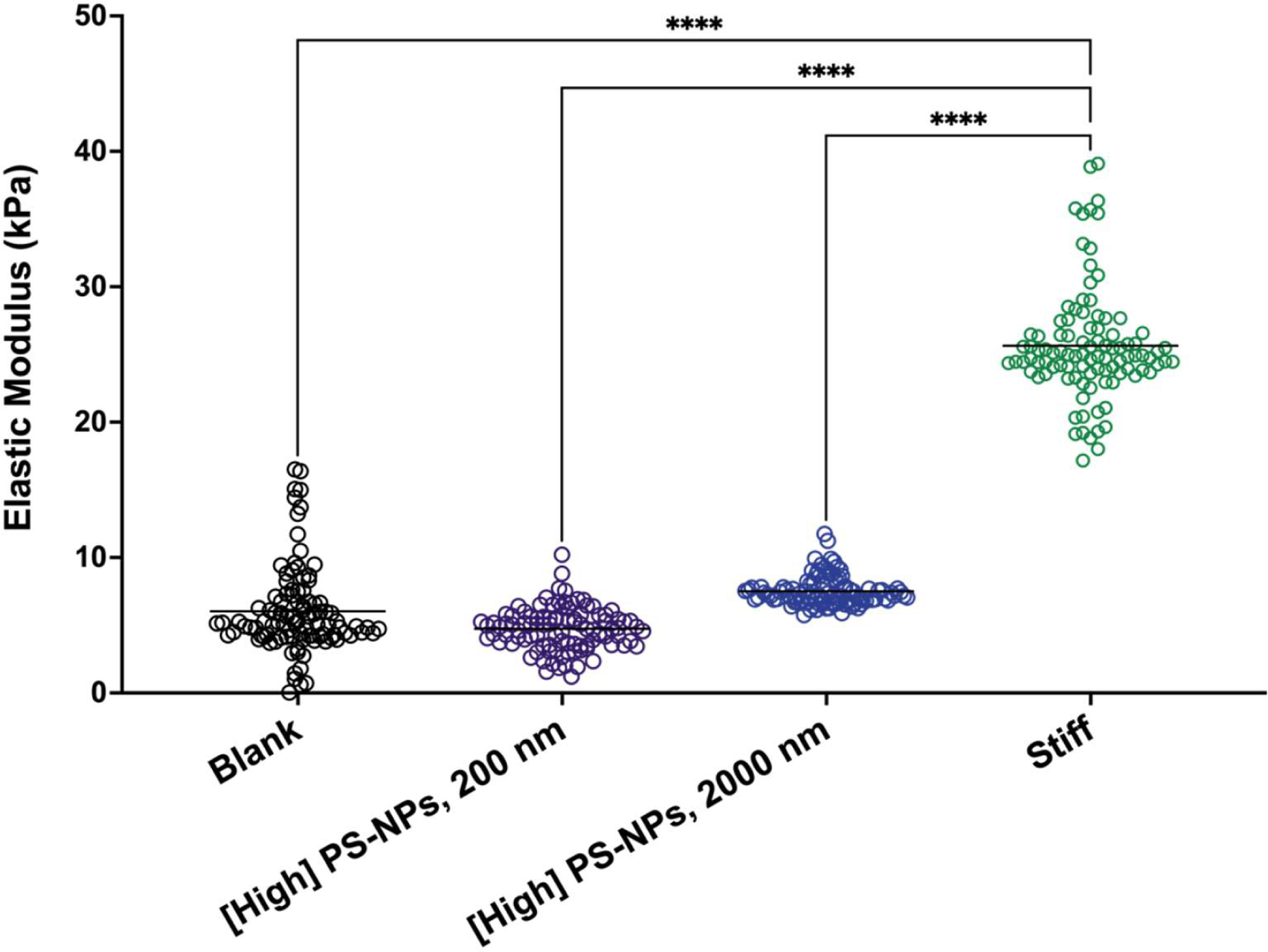
Atomic force microscopy (AFM) characterization of blank and PS-NPs hydrogels. Elastic moduli of blank gels, blank gels with 200 nm PS-NPs, blank gels with 2000 nm PS-NPs, and stiff gels (n=50 measurements). Significance determined via one-way ANOVA (****p<0.0001 denoting significance between hydrogel formulations).

**Supplementary Figure 8:**
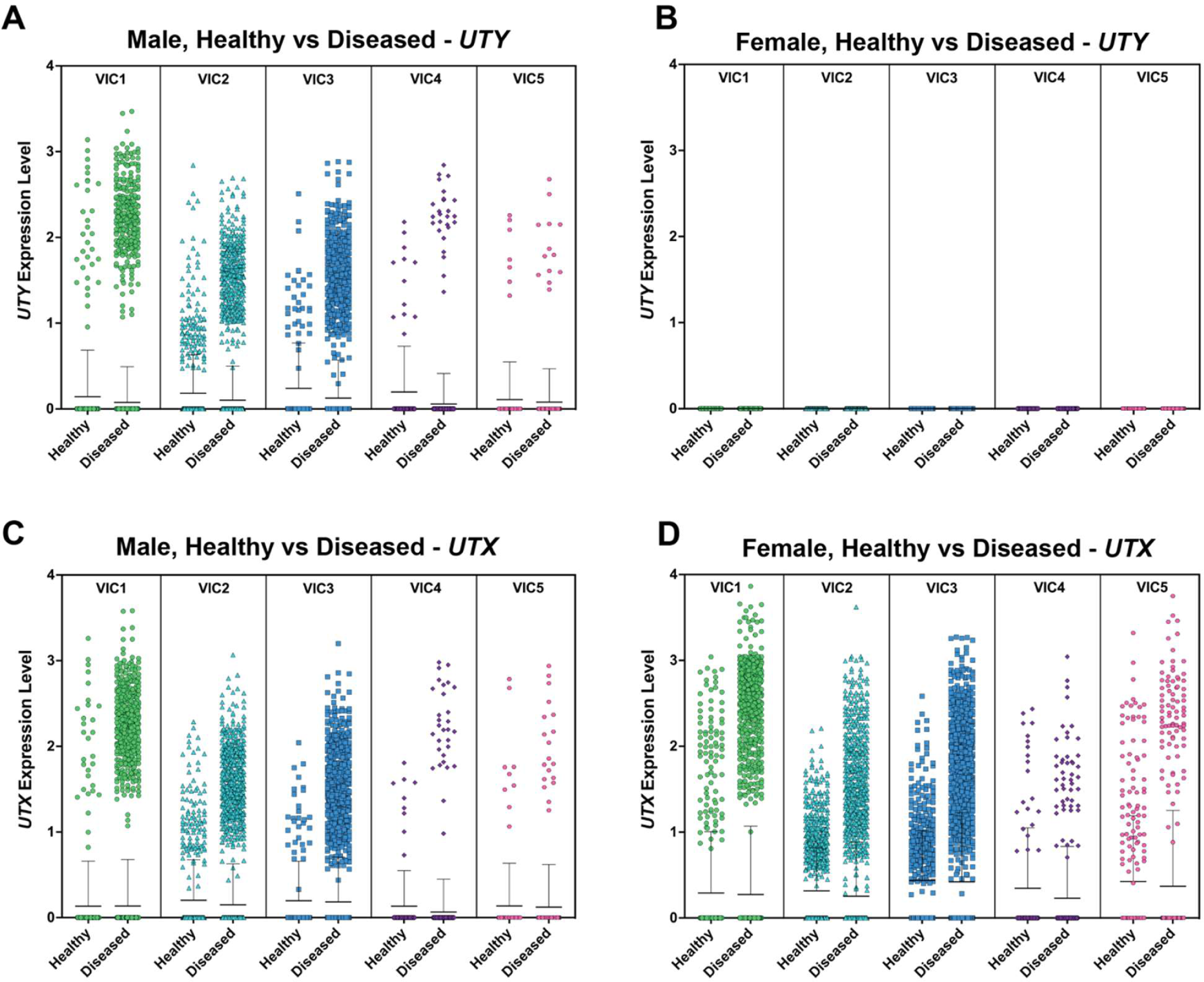
*UTY* and *UTX* mRNA expression in all VIC populations in patient samples. *UTY* relative mRNA expression in VIC populations 1 through 5 in (**A**) males and (**B**) females. *UTX* relative mRNA expression in VIC populations 1 through 5 in (**C**) males and (**D**) females.

**Supplementary Figure 9.**
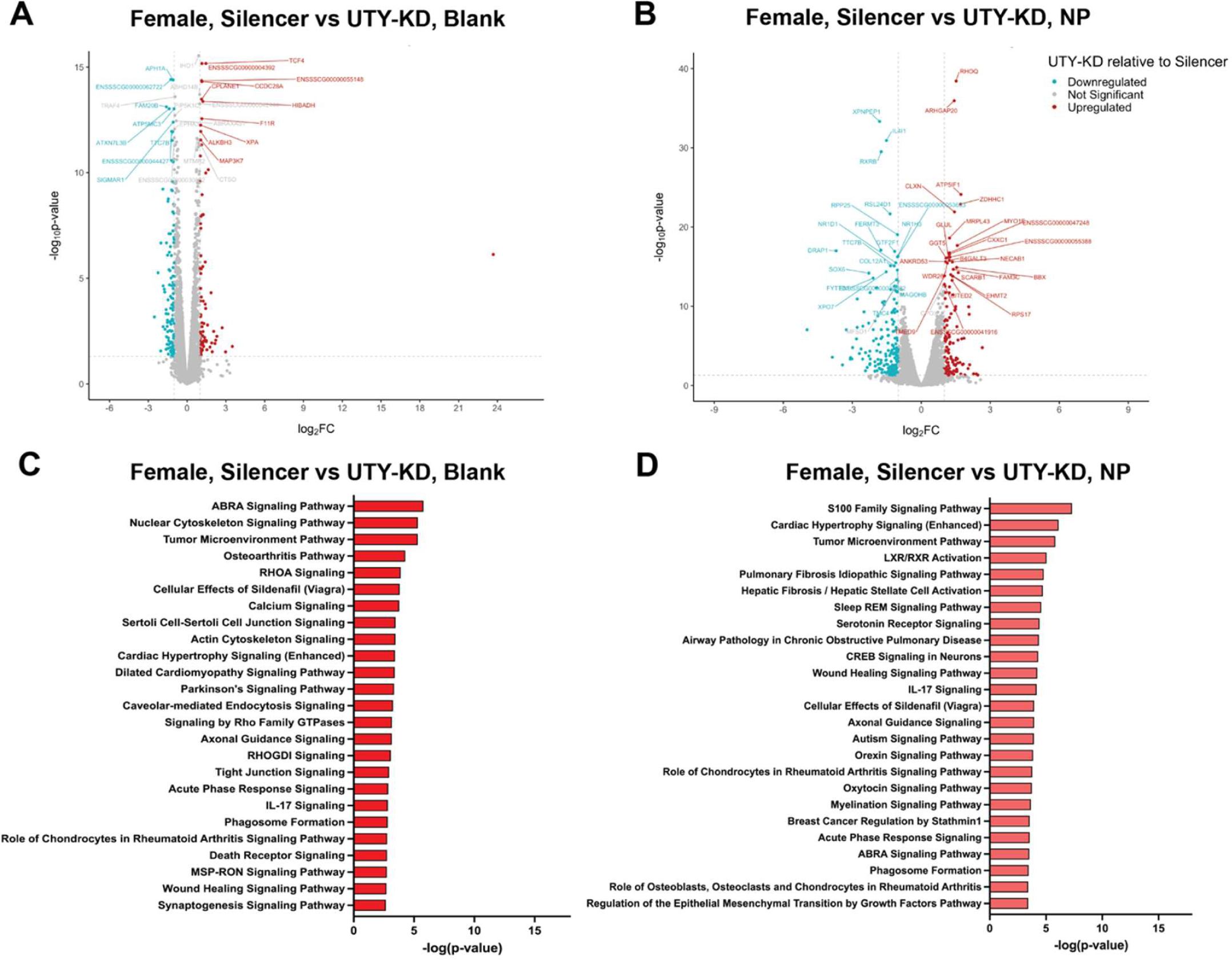
Bulk sequencing and ingenuity pathway analysis in female porcine VICs. Bulk mRNA sequencing shows differentially expressed genes in female VICs (blue = downregulated, red = upregulated, gray = not significant) in (**A**) blank and (**B**) PS-NP hydrogels using DESeq2 (n = 2 biological replicates). Associated pathways from ingenuity pathway analysis in female VICs in (**C**) blank and (**D**) PS-NP hydrogels.

**Supplementary Figure 10:**
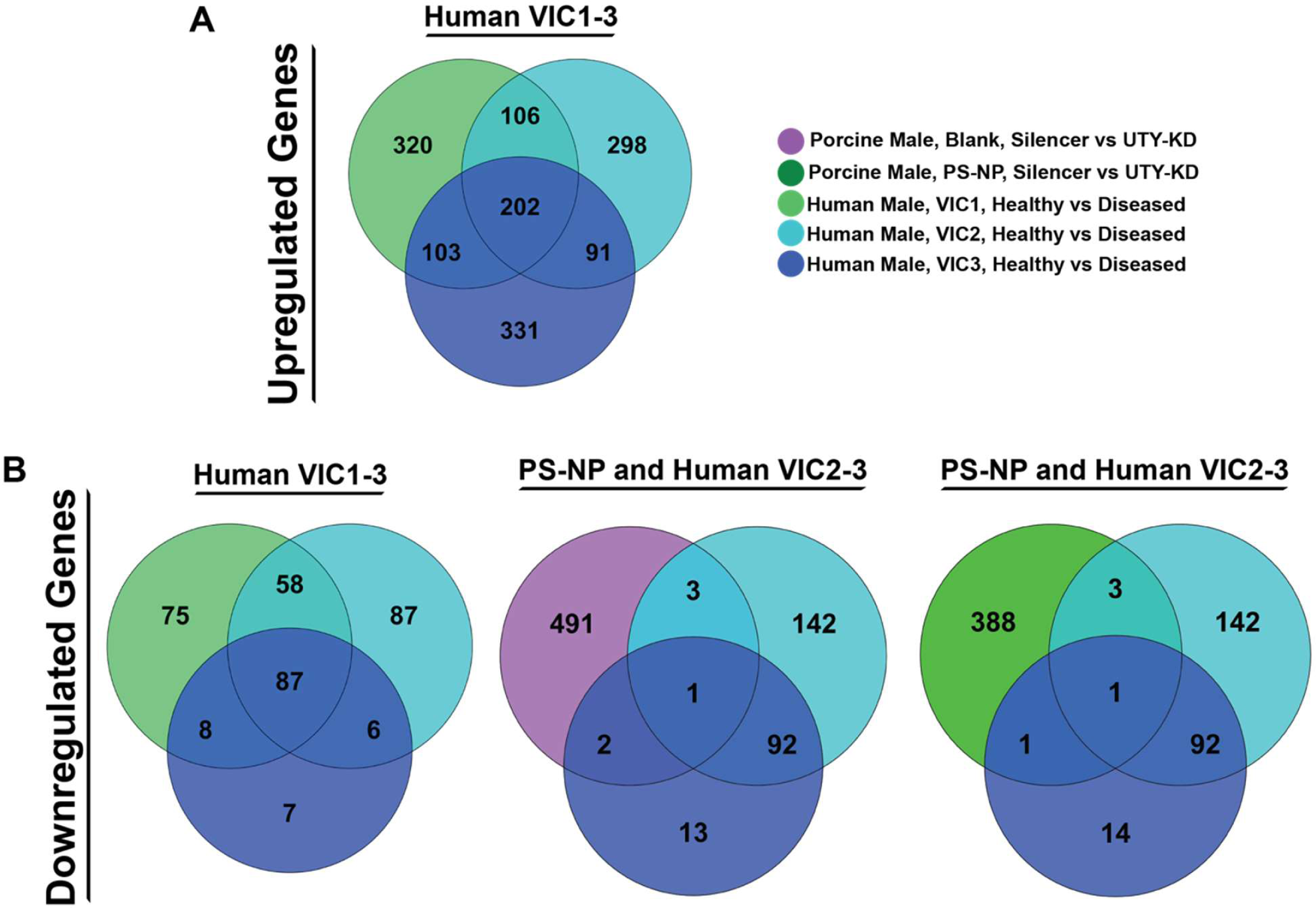
Venn diagrams of common genes across porcine and human sequencing sample comparisons (VIC1-3). (**A**) Venn diagrams of upregulated genes across conditions specified in legend. (**B**) Venn diagrams of downregulated genes across conditions specified in legend.

**Supplementary Figure 11:**
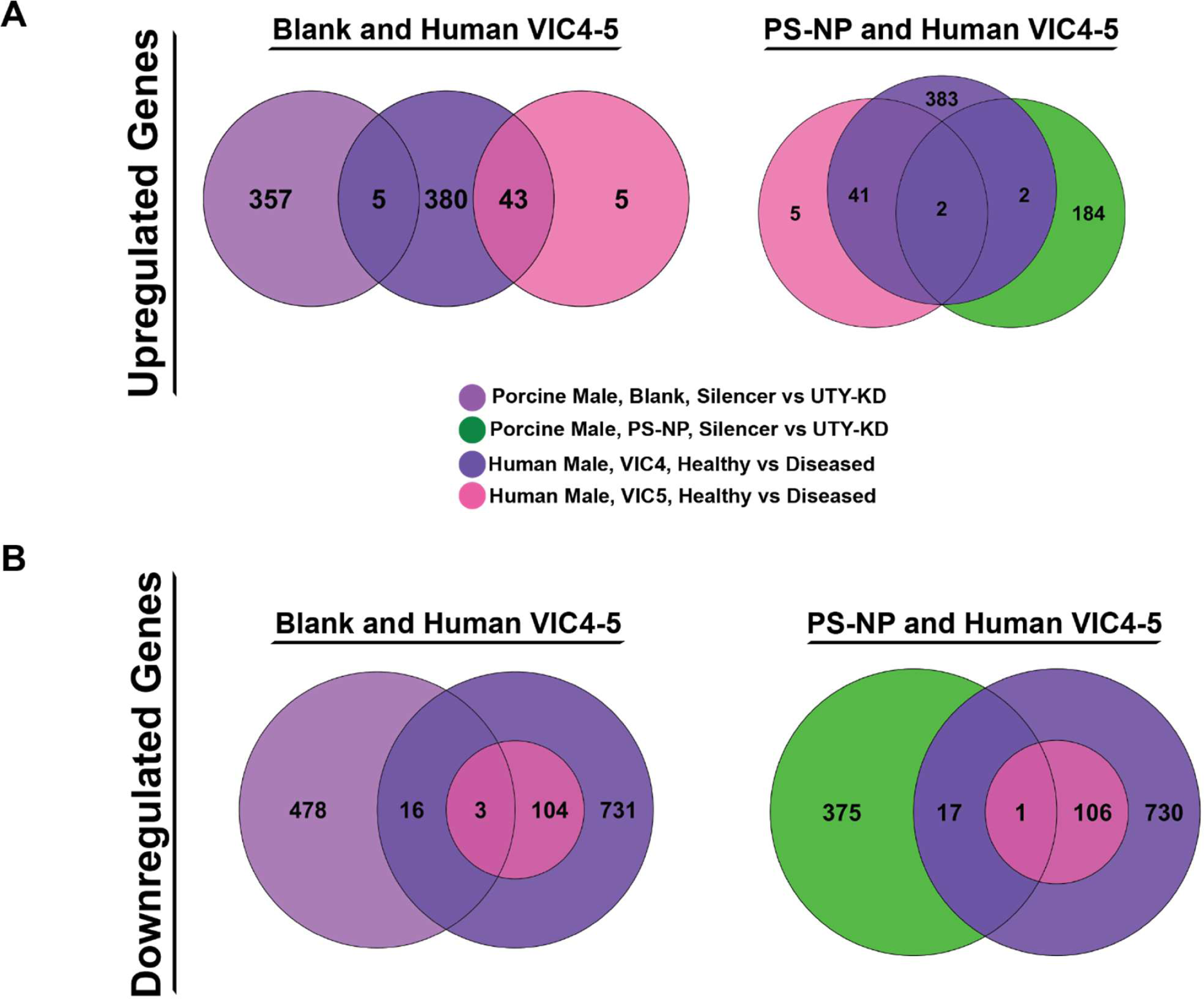
Venn diagrams of common genes across porcine and human sequencing sample comparisons (VIC4-5). (**A**) Venn diagrams of upregulated genes across conditions specified in legend. (**B**) Venn diagrams of downregulated genes across conditions specified in legend.

**Supplementary Figure 12.**
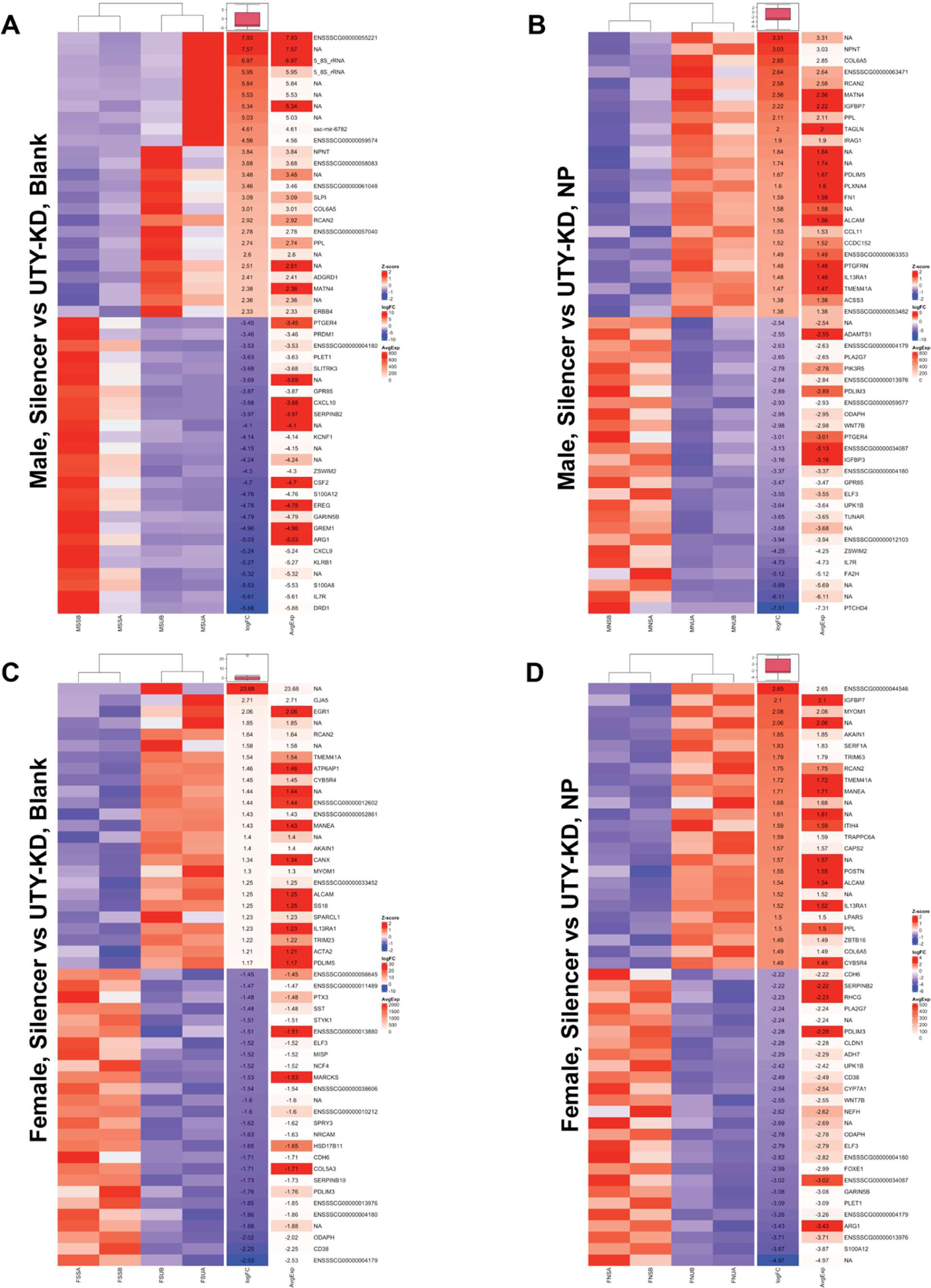
Heatmap of top differentially expressed genes after UTY-knockdown. Top differentially expressed genes (blue = downregulated, red = upregulated) after UTY-knockdown (UTY-KD) for male VICs on (**A**) blank hydrogels, (**B**) male VICs on PS-NP hydrogels, (**C**) female VICs on blank hydrogels, and (**D**) female VICs on PS-NP hydrogels.

**Supplementary Table 1.** Human patient data.

| Sample | Sex | Sample Type | Age | Comorbidities | NCBI GEO Data Accession Number |
| --- | --- | --- | --- | --- | --- |
| Patient 1 | Male | Healthy | 50 | Unknown | PRJNA562645 |
| Patient 2 | Male | CAVD | 50 | Unknown | PRJNA562645 |
| Patient 3 | Male | AVS | 79 | Listed below<sup>a</sup> | GSE273980 |
| Patient 4 | Female | Healthy | 48 | Unknown | PRJNA562645 |
| Patient 5 | Female | CAVD | 52 | Unknown | PRJNA562645 |
| Patient 6 | Female | AVS | 76 | Listed below<sup>b</sup> | GSE273980 |

<sup>a</sup>) Cerebrovascular accident (CVA, Stroke), Congestive heart failure (CHF), Atrial fibrillation (Afib, A-fib, AF), Aortic (valve) stenosis, Chronic obstructive pulmonary disease (COPD), Pulmonary hypertension (pHTN, pulmonary HTN), Chronic kidney disease (CKD), Diabetes - Type II (T2D, T2DM), Hypothyroidism, Gastroesophageal reflux disease (GERD), Anemia, Thrombocytopenia, Coronary artery disease (CAD)
<sup>b</sup>) Atherosclerosis, Angina, Atrial fibrillation (Afib, A-fib, AF), Cardiac pacemaker, Pulmonary hypertension (pHTN, pulmonary HTN), Aortic (valve) stenosis, Chronic kidney disease (CKD), Heart Failure, Chronic obstructive pulmonary disease (COPD), Congestive heart failure (CHF), Coronary artery disease (CAD), Mitral (valve) stenosis, Anxiety disorder, Anemia, Emphysema, Diabetes - Type II (T2D, T2DM, Duration: 5 Years), Myocardial infarction (MI, heart attack), Cataract - bilateral, Arthritis

**Supplementary Table 2.**
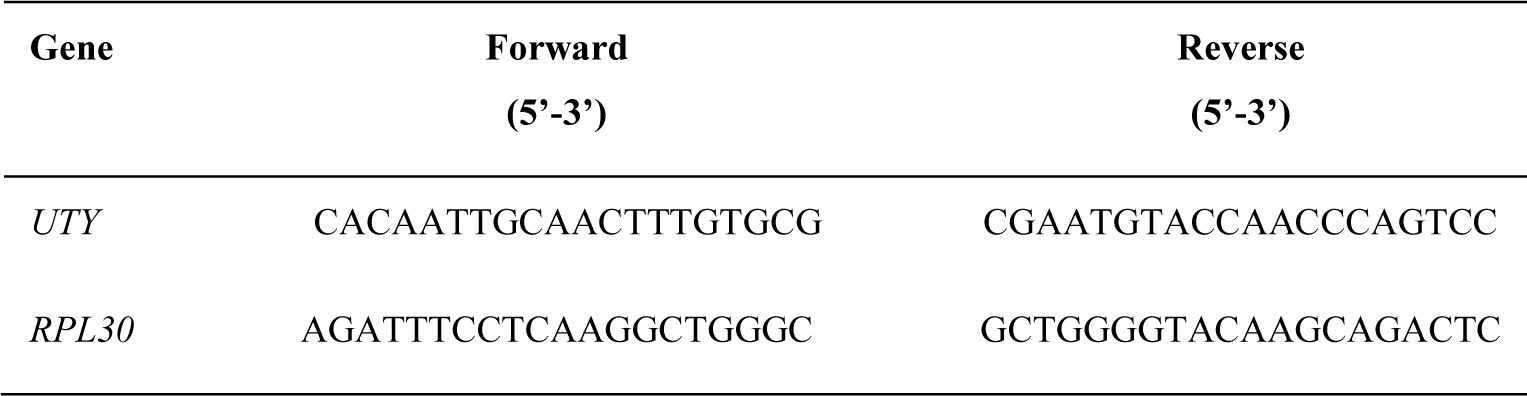
RT-qPCR Porcine Primers.

**Supplementary Table 3.** Bulk RNA sequencing results statistics.

| **Sample** | **Number of Reads** | **% of ≥Q30 Bases** | **Mean Quality Score** | **% of Raw Clusters** | **% of Perfect Index Reads** |
| --- | --- | --- | --- | --- | --- |
| MSS-A | 77,031,106 | 96.1 | 39.20 | 5.57 | 98.02 |
| MSS-B | 70,594,849 | 95.94 | 39.17 | 5.11 | 97.53 |
| MSU-A | 66,197,980 | 94.54 | 38.86 | 4.79 | 97.52 |
| MSU-B | 78,677,637 | 95.91 | 39.17 | 5.69 | 97.77 |
| MNS-A | 75,741,443 | 96.18 | 39.22 | 5.48 | 97.06 |
| MNS-B | 75,188,271 | 96.15 | 39.21 | 5.44 | 98.50 |
| MNU-A | 72,527,065 | 96.21 | 39.23 | 5.25 | 95.60 |
| MNU-B | 71,457,271 | 95.98 | 39.18 | 5.17 | 97.08 |
| FSS-A | 80,582,081 | 96.05 | 39.20 | 5.83 | 98.15 |
| FSS-B | 77,318,130 | 96.12 | 39.21 | 5.59 | 97.76 |
| FSU-A | 73,159,248 | 96.06 | 39.20 | 5.29 | 96.98 |
| FSU-B | 75,757,306 | 96.15 | 39.21 | 5.48 | 97.73 |
| FNS-A | 80,347,740 | 96.12 | 39.21 | 5.81 | 97.90 |
| FNS-B | 69,139,710 | 95.85 | 39.15 | 5.00 | 97.77 |
| FNU-A | 76,303,407 | 96.10 | 39.20 | 5.52 | 97.07 |
| FNU-B | 74,652,434 | 96.08 | 39.20 | 5.40 | 98.17 |

**Supplementary Table 4.** Single cell sequencing results statistics.

| Sample | Number of Reads | Mean Reads per Cell | Valid Barcodes | Q30 Bases in Barcode | Q30 Bases in RNA Read | Q30 Bases in Sample Index | Q30 Bases in UMI |
| --- | --- | --- | --- | --- | --- | --- | --- |
| Patient 3 | 41,797,983 | 6,562 | 96.7% | 96.9% | 96.0% | 96.9% | 97.9% |
| Patient 6 | 39,845,415 | 4,301 | 96.00% | 97.10% | 97.10% | 97.10% | 98.00% |

**Supplementary Table 5.** Single cell sequencing genomic mapping statistics.

| Sample | Reads Mapped Confidently to Transcriptome | Reads Mapped Confidently to Exonic Regions | Reads Mapped Confidently to Intronic Regions | Reads Mapped Confidently to Intergenic Regions | Sequencing Saturation |
| --- | --- | --- | --- | --- | --- |
| Patient 3 | 96.1% | 76.0% | 12.4% | 5.0% | 49.7% |
| Patient 6 | 96.5% | 74.1% | 13.6% | 4.8% | 38.4% |

**Supplementary Table 6.**
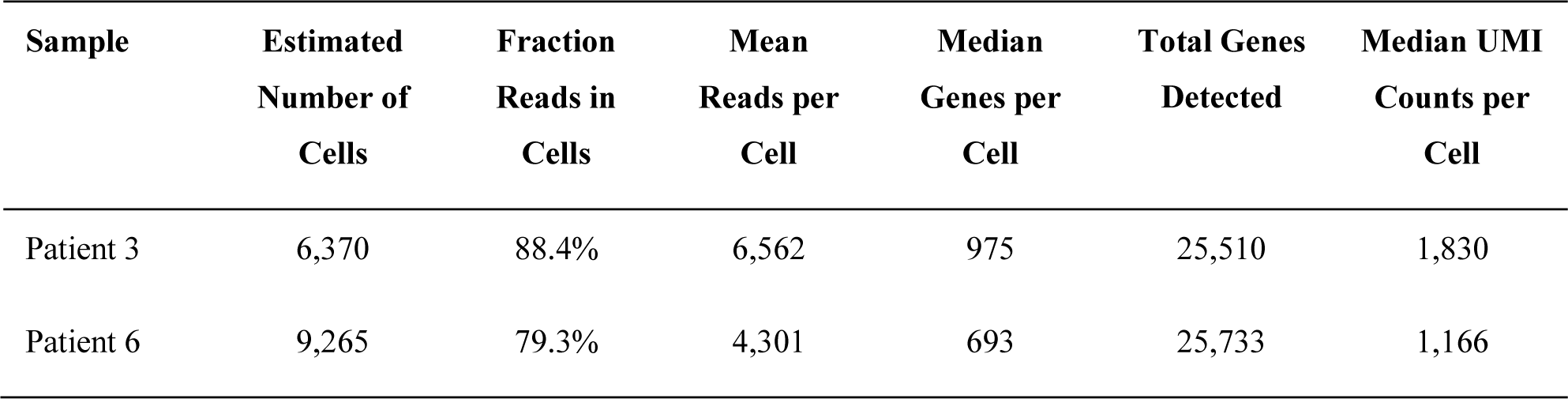
Single cell sequencing gene expression statistics.

**Supplementary Table 7.** Epigenetic modifiers present in porcine and human data.

| Sample | Epigenetic Modifiers |
| --- | --- |
| Porcine, Male, Blank Hydrogel, UTY-KD, Upregulated | TEX10, BRD3, NOC2L, SIRT2, CUL3, NEK9, SMC1A, BRPF3, PRKCB, SP1, SMARCC2, MSL1 |
| Porcine, Male, PS-NP Hydrogel, UTY-KD, Upregulated | SUPT3H, ZNF687, ACTB, ATAD2B, DNMT3A, TAF6 |
| Human, Male, Diseased, VIC1, Upregulated | SUPT3H, JAK2, ARID1B, BRWD3, L3MBTL4, KDM6B, PARG, FTO, SETD2, RAD54L2,CLOCK, PHIP, EYA3, NBN, MGA, NCOA1, ZMYM4, STK4, INO80D, SMARCAD1, NCOA2, MLLT6, ELP2, BTAF1, RNF168, TEX10, TET2, TRRAP, EYA4, TOP2B, MAP3K7, PHC3, ATR, NAT10, KAT6A, TET3, ZNF217, SENP1, TRIM33, KDM5A, ZMYM6, BRD8, CHEK1, APOBEC2 |
| Human, Male, Diseased, VIC2, Upregulated | SUPT3H, DDX21, STK4, ARID1B, KDM6B, USP12, SMCHD1, L3MBTL4, PPP4R3B, PARG, NBN, ERCC6, CHUK, MAP3K7, BTAF1, CDK17, USP3, JAK2, SMARCAD1, BRWD3, HELLS, ALKBH1, TET3, DOT1L, PHF19, CHEK1, APOBEC2, ZNF217, TRIM24, KDM4A, ZNF711, JADE3, SCML2, RAD51 |
| Human, Male, Diseased, VIC3, Upregulated | SUPT3H, DDX21, RNF168, SMARCAD1, ZMYM4, ARID1B, PARG, NBN, L3MBTL4, APOBEC2, TAF3, STK4, RBBP5, EXOSC6, SCMH1, USP36, SSRP1, ATAD2, CDK17, KDM6B, MLLT6, SETD2, ERCC6, USP11, USP15, BRWD3, RAD54L2, TAF1, SHPRH, EHMT1, CHUK, PRDM5, DNMT1, BAZ2A, ATR, INO80, REST, HELLS, CHEK1, BRD3, IKZF1, MAP3K7, SENP3, SETD1A, DZIP3, TYW5, JAK2, JADE3 |

**Supplementary Table 8.**
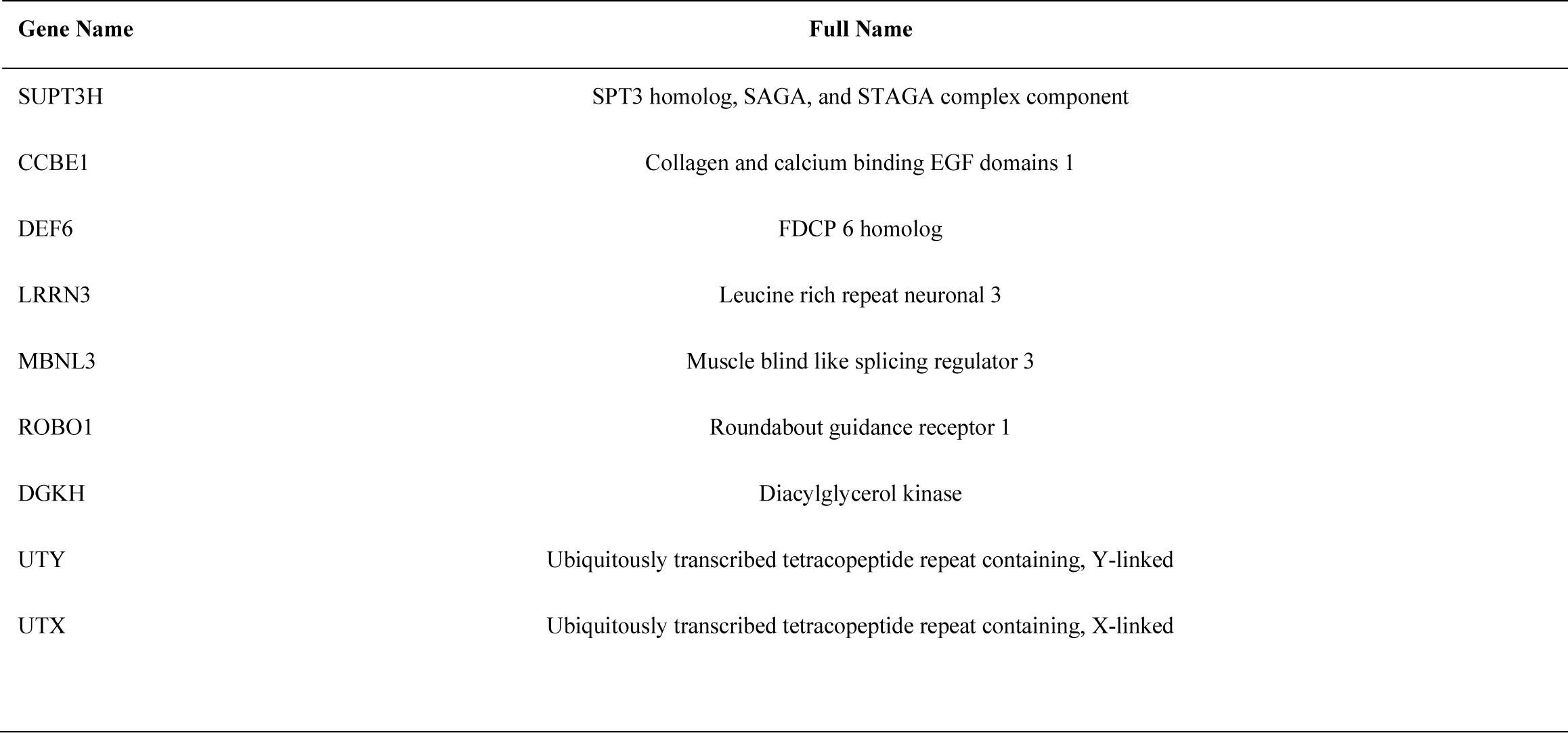
Gene names and abbreviations used.

## References

1. Carabello BA, Paulus WJ. Aortic stenosis. The Lancet. 2009;373(9667):956-966. doi:10.1016/S0140-6736(09)60211-7

2. Raddatz MA, Gonzales HM, Farber-Eger E, Wells QS, Lindman BR, Merryman WD. Characterisation of aortic stenosis severity: a retrospective analysis of echocardiography reports in a clinical laboratory. Open Heart. 2020;7(2):e001331. doi:10.1136/openhrt-2020-001331

3. Myasoedova VA, Ravani AL, Frigerio B, et al. Novel pharmacological targets for calcific aortic valve disease: Prevention and treatments. Pharmacol Res. 2018;136:74–82. doi:10.1016/j.phrs.2018.08.020

4. Simard L, Côté N, Dagenais F, et al. Sex-Related Discordance Between Aortic Valve Calcification and Hemodynamic Severity of Aortic Stenosis | Circulation Research. Circ Res. 120. https://www.ahajournals.org/doi/full/10.1161/CIRCRESAHA.116.309306

5. Gourgas O, Khan K, Schwertani A, Cerruti M. Differences in mineral composition and morphology between men and women in aortic valve calcification. Acta Biomater. 2020;106:342–350. doi:10.1016/j.actbio.2020.02.030

6. Ayoub S, Lee CH, Driesbaugh KH, et al. Regulation of valve interstitial cell homeostasis by mechanical deformation: implications for heart valve disease and surgical repair. J R Soc Interface. 2017;14(135):20170580. doi:10.1098/rsif.2017.0580

7. Goody PR, Hosen MR, Christmann D, et al. Aortic Valve Stenosis. Arterioscler Thromb Vasc Biol. 2020;40(4):885–900. doi:10.1161/ATVBAHA.119.313067

8. Blaser MC, Kraler S, Lüscher TF, Aikawa E. Multi-Omics Approaches to Define Calcific Aortic Valve Disease Pathogenesis. Circ Res. 2021;128(9):1371–1397. doi:10.1161/CIRCRESAHA.120.317979

9. Nagy E, Eriksson P, Yousry M, et al. Valvular osteoclasts in calcification and aortic valve stenosis severity. Int J Cardiol. 2013;168(3):2264–2271. doi:10.1016/j.ijcard.2013.01.207

10. Yu C, Li L, Xie F, et al. LncRNA TUG1 sponges miR-204-5p to promote osteoblast differentiation through upregulating Runx2 in aortic valve calcification. Cardiovasc Res. 2018;114(1):168–179. doi:10.1093/cvr/cvx180

11. Bertazzo S, Gentleman E. Aortic valve calcification: a bone of contention. Eur Heart J. 2017;38(16):1189–1193. doi:10.1093/eurheartj/ehw071

12. Ibrahim M, Schoelermann J, Mustafa K, Cimpan MR. TiO2 nanoparticles disrupt cell adhesion and the architecture of cytoskeletal networks of human osteoblast-like cells in a size dependent manner. J Biomed Mater Res A. 2018;106(10):2582–2593. doi:10.1002/jbm.a.36448

13. Chen JH, Simmons CA, Towler DA. Cell–Matrix Interactions in the Pathobiology of Calcific Aortic Valve Disease. Circ Res. 2011;108(12):1510–1524. doi:10.1161/CIRCRESAHA.110.234237

14. Kodigepalli KM, Thatcher K, West T, et al. Biology and Biomechanics of the Heart Valve Extracellular Matrix. J Cardiovasc Dev Dis. 2020;7(4):57. doi:10.3390/jcdd7040057

15. Barczyk M, Carracedo S, Gullberg D. Integrins. Cell Tissue Res. 2010;339(1):269-280. doi:10.1007/s00441-009-0834-6

16. Takada Y, Ye X, Simon S. The integrins. Genome Biol. 2007;8(5):215. doi:10.1186/gb-2007-8-5-215

17. van der Flier A, Sonnenberg A. Function and interactions of integrins. Cell Tissue Res. 2001;305(3):285-298. doi:10.1007/s004410100417

18. Walker CJ, Crocini C, Ramirez D, et al. Nuclear mechanosensing drives chromatin remodelling in persistently activated fibroblasts. Nat Biomed Eng. 2021;5(12):1485–1499. doi:10.1038/s41551-021-00709-w

19. Walker CJ, Batan D, Bishop CT, et al. Extracellular matrix stiffness controls cardiac valve myofibroblast activation through epigenetic remodeling. Bioeng Transl Med. 2022;7(3):e10394. doi:10.1002/btm2.10394

20. Walker CJ, Schroeder ME, Aguado BA, Anseth KS, Leinwand LA. Matters of the heart: Cellular sex differences. J Mol Cell Cardiol. 2021;160:42–55. doi:10.1016/j.yjmcc.2021.04.010

21. Hartman RJG, Huisman SE, den Ruijter HM. Sex differences in cardiovascular epigenetics—a systematic review. Biol Sex Differ. 2018;9(1):19. doi:10.1186/s13293-018-0180-z

22. Deegan DF, Nigam P, Engel N. Sexual Dimorphism of the Heart: Genetics, Epigenetics, and Development. Front Cardiovasc Med. 2021;8. doi:10.3389/fcvm.2021.668252

23. Wijchers PJ, Festenstein RJ. Epigenetic regulation of autosomal gene expression by sex chromosomes. Trends Genet. 2011;27(4):132–140. doi:10.1016/j.tig.2011.01.004

24. Dobson LE, Fairbairn TA, Plein S, Greenwood JP. Sex Differences in Aortic Stenosis and Outcome Following Surgical and Transcatheter Aortic Valve Replacement. J Womens Health. 2015;24(12):986–995. doi:10.1089/jwh.2014.5158

25. Côté N, Clavel MA. Sex Differences in the Pathophysiology, Diagnosis, and Management of Aortic Stenosis. Cardiol Clin. 2020;38(1):129–138. doi:10.1016/j.ccl.2019.09.008

26. Fleury MA, Annabi MS, Voisine M, et al. Impact of sex and sex hormones on pathophysiology and progression of aortic stenosis in a murine model - Fleury - 2022 - Physiological Reports - Wiley Online Library. Physiol Soc. 10(16). https://physoc.onlinelibrary.wiley.com/doi/full/10.14814/phy2.15433

27. Aguado BA, Walker CJ, Grim JC, et al. Genes That Escape X Chromosome Inactivation Modulate Sex Differences in Valve Myofibroblasts. Circulation. 2022;145(7):513–530. doi:10.1161/CIRCULATIONAHA.121.054108

28. Vogt BJ, Peters DK, Anseth KS, Aguado BA. Inflammatory serum factors from aortic valve stenosis patients modulate sex differences in valvular myofibroblast activation and osteoblast-like differentiation. Biomater Sci. 2022;10(22):6341–6353. doi:10.1039/D2BM00844K

29. James BD, Allen JB. Sex-Specific Response to Combinations of Shear Stress and Substrate Stiffness by Endothelial Cells In Vitro. Adv Healthc Mater. 2021;10(18):2100735. doi:10.1002/adhm.202100735

30. Sánchez Marrero G, Villa-Roel N, Li F, et al. Single-Cell RNA sequencing investigation of female-male differences under PAD conditions. Front Cardiovasc Med. 2023;10. doi:10.3389/fcvm.2023.1251141

31. Nemec S, Kilian KA. Materials control of the epigenetics underlying cell plasticity. Nat Rev Mater. 2021;6(1):69–83. doi:10.1038/s41578-020-00238-z

32. O’Seaghdha CM, Wu H, Yang Q, et al. Meta-Analysis of Genome-Wide Association Studies Identifies Six New Loci for Serum Calcium Concentrations | PLOS Genetics. PLOS Genet. https://journals.plos.org/plosgenetics/article?id=10.1371/journal.pgen.1003796

33. Howles SA, Wiberg A, Goldsworthy M, et al. Genetic variants of calcium and vitamin D metabolism in kidney stone disease. Nat Commun. 2019;10(1):5175. doi:10.1038/s41467-019-13145-x

34. Binder N, Miller C, Yoshida M, et al. Def6 Restrains Osteoclastogenesis and Inflammatory Bone Resorption. J Immunol. 2017;198(9):3436–3447. doi:10.4049/jimmunol.1601716

35. Logan NJ, Camman M, Williams G, Higgins CA. Demethylation of ITGAV accelerates osteogenic differentiation in a blast-induced heterotopic ossification *in vitro* cell culture model. Bone. 2018;117:149–160. doi:10.1016/j.bone.2018.09.008

36. Georgiadis P, Hebels DG, Valavanis I, et al. Omics for prediction of environmental health effects: Blood leukocyte-based cross-omic profiling reliably predicts diseases associated with tobacco smoking. Sci Rep. 2016;6(1):20544. doi:10.1038/srep20544

37. Wang J, Shi A, Lyu J. A comprehensive atlas of epigenetic regulators reveals tissue-specific epigenetic regulation patterns. Epigenetics. 18(1):2139067. doi:10.1080/15592294.2022.2139067

38. Arumugam B, Vishal M, Shreya S, et al. Parathyroid hormone-stimulation of Runx2 during osteoblast differentiation via the regulation of lnc-SUPT3H-1:16 (RUNX2-AS1:32) and miR-6797-5p. Biochimie. 2019;158:43–52. doi:10.1016/j.biochi.2018.12.006

39. Rice SJ, Aubourg G, Sorial AK, et al. Identification of a novel, methylation-dependent, RUNX2 regulatory region associated with osteoarthritis risk. Hum Mol Genet. 2018;27(19):3464–3474. doi:10.1093/hmg/ddy257

40. Barutcu AR, Tai PWL, Wu H, et al. The bone-specific Runx2-P1 promoter displays conserved three-dimensional chromatin structure with the syntenic Supt3h promoter. Nucleic Acids Res. 2014;42(16):10360–10372. doi:10.1093/nar/gku712

41. Mas-Peiro S, Abplanalp WT, Rasper T, et al. Mosaic loss of Y chromosome in monocytes is associated with lower survival after transcatheter aortic valve replacement. Eur Heart J. 2023;44(21):1943–1952. doi:10.1093/eurheartj/ehad093

42. Sano S, Horitani K, Ogawa H, et al. Hematopoietic loss of Y chromosome leads to cardiac fibrosis and heart failure mortality. Science. 2022;377(6603):292-297. doi:10.1126/science.abn3100

43. Thompson DJ, Genovese G, Halvardson J, et al. Genetic predisposition to mosaic Y chromosome loss in blood. Nature. 2019;575(7784):652–657. doi:10.1038/s41586-019-1765-3

44. Cunningham CM, Li M, Ruffenach G, et al. Y-Chromosome Gene, Uty, Protects Against Pulmonary Hypertension by Reducing Proinflammatory Chemokines | American Journal of Respiratory and Critical Care Medicine. DOI: 10.1164/rccm.202110-2309OC

45. Puperi DS, Balaoing LR, O’Connell RW, West JL, Grande-Allen KJ. 3-Dimensional spatially organized PEG-based hydrogels for an aortic valve co-culture model. Biomaterials. 2015;67:354–364. doi:10.1016/j.biomaterials.2015.07.039

46. Puperi DS, Kishan A, Punske ZE, et al. Electrospun Polyurethane and Hydrogel Composite Scaffolds as Biomechanical Mimics for Aortic Valve Tissue Engineering. ACS Biomater Sci Eng. 2016;2(9):1546–1558. doi:10.1021/acsbiomaterials.6b00309

47. Porras AM, Westlund JA, Evans AD, Masters KS. Creation of disease-inspired biomaterial environments to mimic pathological events in early calcific aortic valve disease. Proc Natl Acad Sci. 2018;115(3):E363–E371. doi:10.1073/pnas.1704637115

48. Scott AJ, Simon LR, Hutson HN, Porras AM, Masters KS. Engineering the aortic valve extracellular matrix through stages of development, aging, and disease. J Mol Cell Cardiol. 2021;161:1–8. doi:10.1016/j.yjmcc.2021.07.009

49. Floy ME, Shabnam F, Givens SE, et al. Identifying molecular and functional similarities and differences between human primary cardiac valve interstitial cells and ventricular fibroblasts. Front Bioeng Biotechnol. 2023;11. Accessed July 2, 2023. https://www.frontiersin.org/articles/10.3389/fbioe.2023.1102487

50. Driscoll K, Cruz AD, Butcher JT. Inflammatory and Biomechanical Drivers of Endothelial-Interstitial Interactions in Calcific Aortic Valve Disease. Circ Res. 2021;128(9):1344–1370. doi:10.1161/CIRCRESAHA.121.318011

51. Grim JC, Aguado BA, Vogt BJ, et al. Secreted Factors From Proinflammatory Macrophages Promote an Osteoblast-Like Phenotype in Valvular Interstitial Cells. Arterioscler Thromb Vasc Biol. 2020;40(11):e296–e308. doi:10.1161/ATVBAHA.120.315261

52. Zhang J, Chen Y, Gao M, et al. Silver Nanoparticles Compromise Female Embryonic Stem Cell Differentiation through Disturbing X Chromosome Inactivation | ACS Nano. https://pubs.acs.org/doi/full/10.1021/acsnano.8b08604

53. Weivoda MM, Chew CK, Monroe DG, et al. Identification of osteoclast-osteoblast coupling factors in humans reveals links between bone and energy metabolism. Nat Commun. 2020;11(1):87. doi:10.1038/s41467-019-14003-6

54. Montañés-Agudo P, Pinto YM, Creemers EE. Splicing factors in the heart: Uncovering shared and unique targets. J Mol Cell Cardiol. 2023;179:72–79. doi:10.1016/j.yjmcc.2023.04.003

55. Arnold AP, Cassis LA, Eghbali M, Reue K, Sandberg K. Sex Hormones and Sex Chromosomes Cause Sex Differences in the Development of Cardiovascular Diseases. Arterioscler Thromb Vasc Biol. 2017;37(5):746–756. doi:10.1161/ATVBAHA.116.307301

56. Link JC, Chen X, Arnold AP, Reue K. Metabolic impact of sex chromosomes. Adipocyte. 2013;2(2):74–79. doi:10.4161/adip.23320

57. Burgoyne PS, Arnold AP. A primer on the use of mouse models for identifying direct sex chromosome effects that cause sex differences in non-gonadal tissues. Biol Sex Differ. 2016;7(1):68. doi:10.1186/s13293-016-0115-5

58. Richards JM, Kunitake JAMR, Hunt HB, et al. Crystallinity of hydroxyapatite drives myofibroblastic activation and calcification in aortic valves. Acta Biomater. 2018;71:24–36. doi:10.1016/j.actbio.2018.02.024

59. Vélez NEF, Gorashi RM, Aguado BA. Chemical and molecular tools to probe biological sex differences at multiple length scales. J Mater Chem B. 2022;10(37):7089–7098. doi:10.1039/D2TB00871H

60. Fogg K, Tseng NH, Peyton SR, et al. Roadmap on biomaterials for women’s health. J Phys Mater. 2022;6(1):012501. doi:10.1088/2515-7639/ac90ee

61. Aguado BA, Schuetze KB, Grim JC, et al. Transcatheter aortic valve replacements alter circulating serum factors to mediate myofibroblast deactivation. Sci Transl Med. 2019;11(509):eaav3233. doi:10.1126/scitranslmed.aav3233

62. Xu K, Xie S, Huang Y, et al. Cell-Type Transcriptome Atlas of Human Aortic Valves Reveal Cell Heterogeneity and Endothelial to Mesenchymal Transition Involved in Calcific Aortic Valve Disease. Arterioscler Thromb Vasc Biol. 2020;40(12):2910–2921. doi:10.1161/ATVBAHA.120.314789

63. Satija R, Farrell JA, Gennert D, Schier AF, Regev A. Spatial reconstruction of single-cell gene expression data. Nat Biotechnol. 2015;33(5):495–502. doi:10.1038/nbt.3192

64. Butler A, Hoffman P, Smibert P, Papalexi E, Satija R. Integrating single-cell transcriptomic data across different conditions, technologies, and species. Nat Biotechnol. 2018;36(5):411–420. doi:10.1038/nbt.4096

65. Stuart T, Butler A, Hoffman P, et al. Comprehensive Integration of Single-Cell Data. Cell. 2019;177(7):1888–1902.e21. doi:10.1016/j.cell.2019.05.031

66. Hao Y, Hao S, Andersen-Nissen E, et al. Integrated analysis of multimodal single-cell data. Cell. 2021;184(13):3573–3587.e29. doi:10.1016/j.cell.2021.04.048

67. Hao Y, Stuart T, Kowalski MH, et al. Dictionary learning for integrative, multimodal and scalable single-cell analysis. Nat Biotechnol. 2024;42(2):293–304. doi:10.1038/s41587-023-01767-y

